# DNA origami directed virus capsid polymorphism

**DOI:** 10.1101/2022.11.07.515152

**Authors:** Iris Seitz, Esa-Pekka Kumpula, Eduardo Anaya-Plaza, Jeroen J. L. M. Cornelissen, Veikko Linko, Juha T. Huiskonen, Mauri A. Kostiainen

## Abstract

Most known viruses protect their genome by encapsulating it inside a protein capsid. Viral capsids can adopt various geometries, most iconically characterized by icosahedral or helical symmetries. The assembly process of native capsids is highly cooperative and governed by the protein geometry, protein-protein as well as protein-nucleic acid interactions. Importantly, the absolute control over the size and shape of virus capsids would have imminent advantages in the development of new vaccines and delivery systems. However, tools to direct the assembly process in a programmable manner are exceedingly elusive or strictly limited to specific structures. Here, we introduce a modular approach by demonstrating DNA origami directed polymorphism of single protein subunit capsids. We achieve control over the capsid shape, size, and topology by employing user-defined DNA origami nanostructures as binding and assembly platforms for the capsid proteins. Binding assays and single-particle cryo-electron microscopy reconstruction show that the DNA origami nanoshapes are efficiently encapsulated within the capsid. Further, we observe that helical arrangement of hexameric capsomers is the preferred mode of packing, while a negative curvature of the origami structure is not well tolerated. The capsid proteins assemble on DNA origami in single or double layer configurations depending on the applied stoichiometry. In addition, the obtained viral capsid coatings are able to efficiently shield the encapsulated DNA origami from nuclease degradation and prevent the structures from aggregation. Therefore, these findings may in addition find direct implementations in DNA nanotechnology-based bioengineering by paving the way for the next-generation cargo protection and targeting strategies.

---

Protein cages can be prepared by *de novo* design, engineering of existing proteins or isolated from nature. Design examples of non-native systems include metal-coordinated cages [1, 2] as well as single- and two-component icosahedral architectures [3, 4]. Such systems have proven to be effective in for example immunogen display and potent vaccine design [5]. On the other hand, native virus capsids have unique assembly properties, making them popular building blocks within nanobioengineering [6–10]. A polymorphic behavior has been observed upon *in vitro* reassembly for several virus types [11–14], however directing the polymorphism in a modular user defined way remains challenging [15, 16].

The spherical cowpea chlorotic mottle virus (CCMV, diameter d = 28 nm) is one of the most studied viruses. Its 180 CP copies are arranged into 20 hexamers and 12 pentamers, resulting in a quasi-icosahedral *T* = 3 symmetry [17]. Its reversible assembly process is well characterized and highly dependent on environmental conditions like pH and ionic strength, resulting in the formation of hexagonal sheets, empty spheres and tubes [18–21]. While the control over spherical assemblies has been demonstrated by the encapsulation of both organic and inorganic materials [22–24], the formation of other assemblies that deviate from the native icosahedral symmetry are not possible to achieve in a modular way.

Here, we utilize DNA origami templates [25] to obtain a precise control of the virus capsid assembly’s size and shape. To this end, a long, single-stranded DNA scaffold strand is assembled into well-defined two- and three-dimensional structures [26, 27] through programmable hybridization with short, singlestranded staple sequences [25, 28]. First we study directing the assembly of CCMV CPs using a 6-helix bundle (6HB) DNA origami (Fig. 1). 6HB fits inside the previously described, hollow tubes [18] (considering also the thickness of the CP layer [17]), and additionally, it closely mimicks the packaging of DNA observed in naturally occurring viral systems [29]. By optimizing the ratio between DNA origami and CPs, complexes with multiple protein layers were obtained and further characterized using single-particle cryo-electron microscopy (cryo-EM) reconstruction. This approach allows not only the encapsulation of non-linear and non-tubular structures, but it also presents a versatile technique for protecting DNA against nuclease degradation, thus making it attractive for vaccine and nucleic acid delivery vector development.

**Fig. 1.**
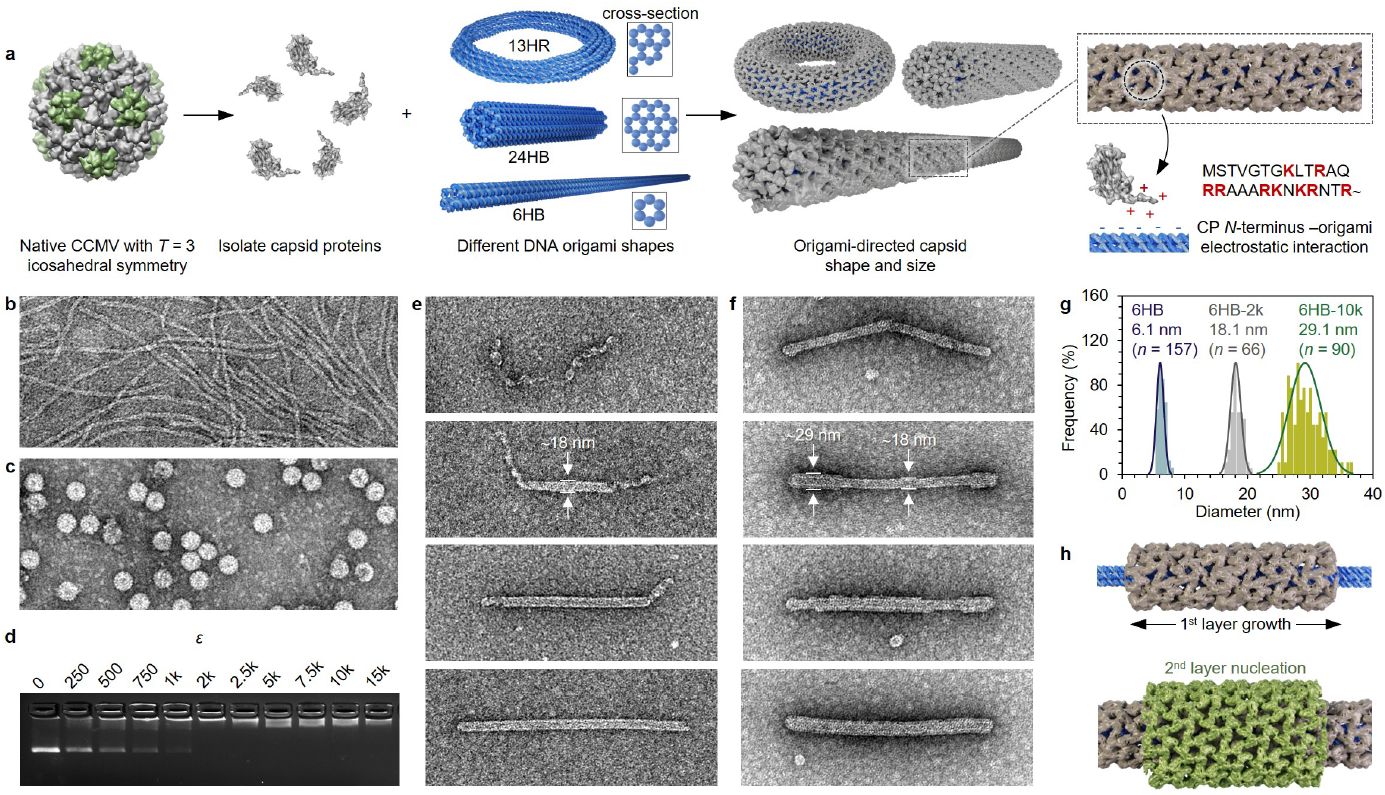
Formation of capsid-coated DNA origami structures. **a**, CPs are isolated from native CCMV (left) and complexed with different DNA origami shapes resulting in a coating (middle) whose properties are determined by the origami structure. The assembly is driven by electrostatic (positively charged aas in the *N*-terminus marked in red) and protein-protein interactions (right). **b**, Negative-stain TEM image of plain 6HB structures. **c**, Negative-stain TEM image of native CCMV particles. **d**, Electrophoretic mobility shift assay (EMSA) shows a mobility decrease of 6HB upon complexation with CPs when increasing *ε*. **e**, Development of a single layer of CPs on the 6HB origami template using *ε* ≤ 2k. **f**, Subsequent development of a second CP layer on top of **e**. **g**, Observed diameter distributions for plain 6HB (blue) and 6HB complexed at *ε* = 2k (grey) or at *ε* = 10k (green). **h**, Schematic presenting the growth of each CP layer based on initial nucleation of the CPs on the DNA origami (top) and a protein layer (bottom). The image width of all TEM images corresponds to 500 nm.

We have used four different DNA origami structures to investigate the properties and possible geometrical limitations that govern the assembly of CCMV CPs. Both 6HB and 24-helix bundle (24HB) are cylindrical structures with diameters of 6 nm and 14 nm, respectively. 13-helix ring (13HR) is a toroidal structure with an outer diameter of 66 nm and a thickness of 9 nm, and 60-helix bundle (60HB) is a brick-like object with dimensions of 20 nm × 20 nm × 31 nm (Supplementary Note S1). The complexation process is driven by protein-protein as well as electrostatic interactions between the negatively charged phosphate backbone of the DNA origami and the positively charged amino acid (aa) residues (+9) in the 26-residual arginine-rich N-terminal region of the protein (190 aa, ~20 kDa) (Fig. 1a).

After characterizing the starting materials, 6HB (*d*_avg_ = 6.1±0.6 nm, s.d.) and intact CCMV (*d*_avg_ = 26.8±1.0 nm, s.d.) (Fig. 1b,c), with negative-stain transmission electron microscopy (TEM), the complexation of isolated CPs and 6HB was performed at physiological conditions (pH 7.3, 150 mM NaCl) using the CPs in excess. The excess, *ε*, is defined as the molar ratio between the protein and the DNA origami (c_CCMV_/c_origami_). The interaction of the components was studied initially based on the change in electrophoretic mobility of the origami during agarose gel electrophoresis (AGE). With increasing *ε*, the 6HB leading band vanishes gradually, while another band appears (Fig. 1d), which corresponds to the well-known fast assembly behavior of CCMV CPs with low cooperativity [30]. According to AGE, this process is completed at ca. *ε* = 2k. The decrease in electrophoretic mobility indicates a significantly less negative surface charge of the origami upon complexation. Negative-stain TEM images at *ε* = 2k reveal single 6HB structures with increased diameter due to a highly ordered protein shell (Fig. 1e, bottom), whereas samples with *ε* < 2k suggest the development of the protein layer from several nucleation sites along the origami structure. When using *ε* > 2k, assembly of free CPs on top of the first protein layer is observed, until a (nearly) complete second layer has been formed at *ε* = 10k (Fig. 1f). Statistical analysis of the diameter of the complexed structures with increasing *ε* (Fig. 1g) shows a clear change from 6.1±0.6 nm (s.d.) (plain 6HB, blue) to 18.1±0.9 nm (s.d.) (6HB-2k, single layer coating, grey) and further to 29.1±2.5 nm (s.d.) (6HB-10k, double layer coating, green). It is notable that the distribution for 6HB with a double layer protein shell is wider, which might be explained by incomplete protein layers. Since no protein multilayers were observed for *ε* < 2k we hypothesize that a fully developed first layer of proteins is required to facilitate the formation of the second one (Fig. 1h).

In general, CCMV assembly has been described to be initiated by CP dimers forming pentamers. It is followed by the addition of dimeric subunits, resulting in pentameric and hexameric capsomeres with curvature [31, 32]. For the assembly of larger icosahedral assemblies, nucleation has been suggested [31], followed by an elongation phase [33]. However, in the absence of RNA, the assembly based on dimers forming hexameric morphological units as nucleation sites has been proposed [17]. In case of low ionic strength, implying stronger protein-DNA compared to protein-protein interactions, *en masse* mechanisms have been suggested for the assembly. Thereby, the protein shell is formed by cooperative rearrangements after initial disordered binding of the CPs to nucleic acids [34, 35]. We expect our protein shells to nucleate into a hexameric lattice. Based on the assembly of brome mosaic virus (BMV, closely related to CCMV) on spherocylindrical substrates [13, 36] nucleation should occur preferentially along the cylindrical region of 6HB. In contrast, nucleation might be uniformly distributed along the entire structure of 24HB due to the difference in the ratio between the spontaneous curvature radius of CPs and the radius of the origami structures.

The geometrical properties of the observed single- and double-layered complexed structures with respect to the outer diameter of the tubes are consistent with previously reported tubes [18, 21, 37, 38]. The length distribution, in contrast, is remarkably different. While the controllability over the assemblies’ length was previously low [37, 39], here, a narrow length distribution of the complexed structures is obtained, suggesting the encapsulation of exactly one 6HB origami per complexed structure.

To confirm the encapsulation of DNA and for detailed structural characterization, single-particle reconstruction was performed based on cryo-EM (Supplementary Notes S2,3). To this end, particles along the filaments from the recorded images (Fig. 2a,d and Supplementary Fig. S2a,b for 6HB-2k and 6HB-10k complexes, respectively) were continuously picked and 2D classified (Fig. 2b,e). 3D refinement of *ab initio* 3D models, obtained from the recorded images only, results in helical structures with 4.3 Å resolution (Fig. 2c, left; Supplementary Fig. S3a) for the first and 7 Å resolution (Fig. 2f; Supplementary Fig. S3c,d) for the second protein layer. The protein tubes are assembled of hexamers (Fig. 2c, top right), similar to tubes formed by *in vitro* assembled human immunodeficiency virus (HIV) [12]. For the atomic model (Fig. 2c, bottom right), CCMV CPs (PDB:1cwp) were flexibly fitted to the cryo-EM density maps.

**Fig. 2.**
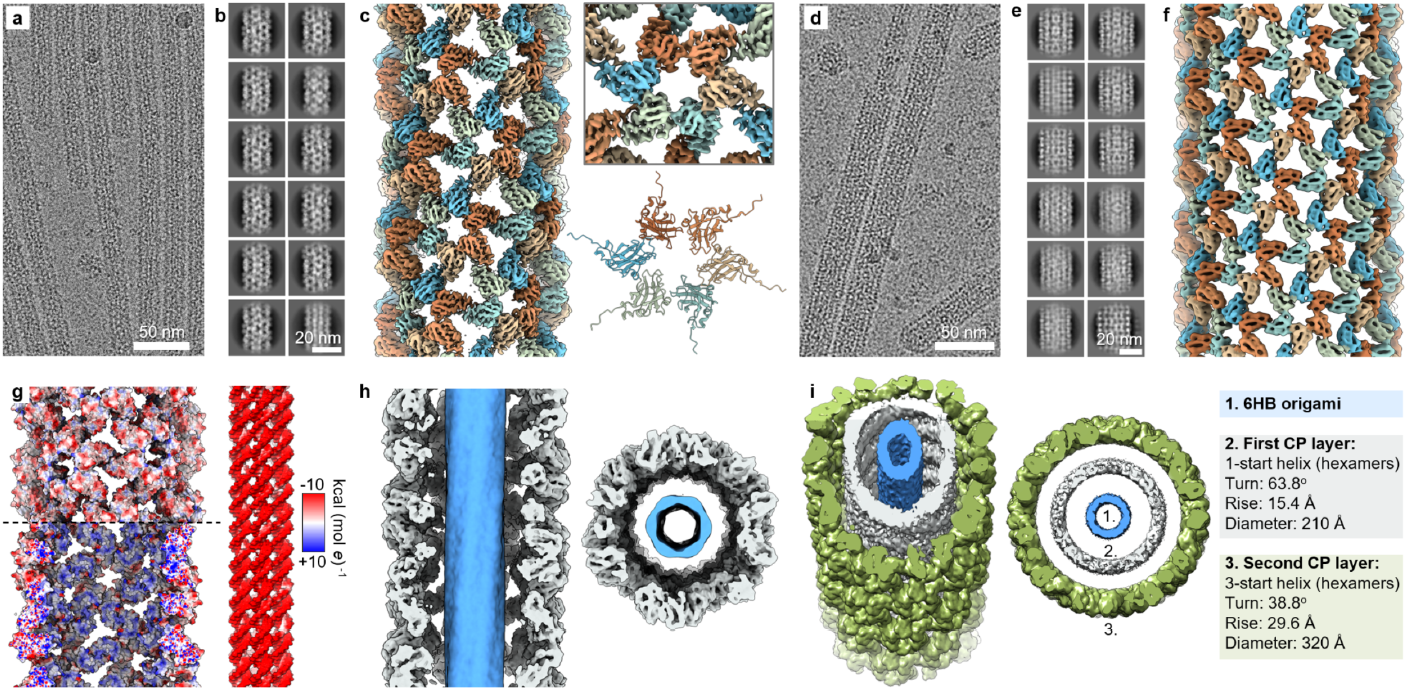
Single-particle reconstruction of complexed 6HB structures using cryo-EM. **a**, Representative micrograph image of 6HB coated with a single protein layer (6HB-2k). **b**, Selected 2D class averages for 6HB-2k. **c**, Cryo-EM density of the tube (left) and a selected hexamer (top right). In an atomic model (bottom right) the CP (PDB:1cwp) was flexibly fitted to the EM density. **d**, Representative image of the double-layered filaments formed by 6HB and the CPs (6HB-10k). **e**, Selected 2D class averages for 6HB-10k. **f**, Cryo-EM density of the outer layer of a double-layered tube. **g**, Electrostatic potential surface suggests a positive potential for the DNA adjacent surface of the inner protein shell (bottom left) while both the outward-facing surface (top left) and the DNA origami (right) possess negative potentials. **h**, Cross-section and top view of 6HB-2k complexes showing the DNA origami in blue. **i**, A 3D model of the assembled double layer structure shows the different symmetry of the two layers assembling on 6HB origami (1). While the inner layer (2) has a 1-start helix symmetry, the outer layer (3) is defined by a 3-start helix.

As expected, the encapsulated DNA origami has its negative charge evenly distributed (Fig. 2g, right). The adjacent protein shell displays a positive electrostatic potential surface caused by the *N*-termini facing towards the origami structure (Fig. 2g, bottom left). For the outer protein surface, in contrast, the negative electrostatic potential is predominant (Fig. 2g, top left; Supplementary Fig. S4d), hence allowing the formation of protein multilayer complexes. We were not able to detect a specific physical contact point between the protein layers, nor between DNA and protein, suggesting unspecific electrostatic as well as protein-protein interactions as the driving force for the assembly.

The density which is clearly visible in the lumen of the protein tubes (Fig. 2h), corresponds to the encapsulated DNA origami, building the core of the assembly. Analysis of the protein layers shows a clear difference in the helical symmetry (Fig, 2i). While the first layer is characterized by a 1-start helix with a turn of 63.8° and a rise of 15.4 Å (Supplementary Fig. S4a), the second layer has a 3-start helical symmetry with a turn of 38.8° and a 29.6 Å rise (Supplementary Fig. S4c,d).

Apart from the increased diameter of the tubular, complexed structures, the helical symmetry is in line with the properties of empty tubes observed by Bancroft *et al*. [18]. CCMV is known to adapt to two distinct conformational states depending on the pH of the surrounding solution [18]. Upon increase of the pH from acidic to neutral and in the absence of metal ions, the capsid transforms from the native state into a swollen state. To this end, the morphology of both hexameric and pentameric units stays unchanged, however, the distance between the units increases by ~5 Å, most likely due to electrostatic repulsion. It is in particular caused by three acidic aa residues at close proximity (most likely with an abnormally high pK_a_ value) which are involved in the metal binding [17]. The cryo-EM densities of CPs encapulating 6HB (Fig. 2, Supplementary Fig. S3) clearly show large holes between the hexameric units, resembling the swollen state which is expected according to the used solution conditions.

The cryo-EM images contained furthermore 3904 ends of the complexed structures, allowing us to subsequently model the cap of the tubes (Fig. 3, Supplementary Note S4) for the first CP layer. The cap is found to consist of six pentamers and one hexamer to create the curvature. Hexamer H0 (Fig. 3b, black) still follows the helical symmetry, while hexamer H1 (blue) is tilted inwards, most likely to be in contact with the pentamers (Fig. 3b-d). The pentamers (P1-5) are arranged along the tubular axis and are responsible for creating the curvature of the cap. Finally, the cap is sealed with a sixth pentamer (P6, Fig. 3d, yellow). Since the diameter of the first protein layer matches with *T* = 1 symmetry, and following from the helical geometry, six pentamers per tube end are expected to close the hexameric lattice which is in line with our volume data.

**Fig. 3.**
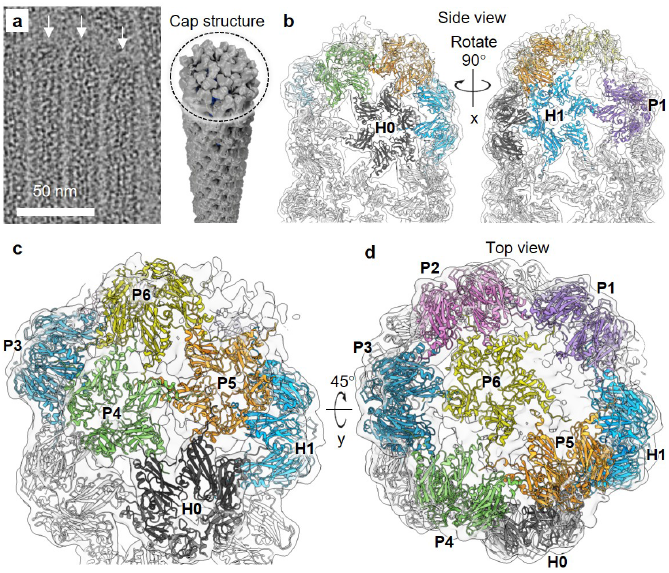
Single-particle reconstruction of the cap structure. **a**, Cryo-EM image of 6HB-2k tubes with caps indicated by white arrows (left) and schematic model of the cap structure (right). **b-d**, Reconstruction of the cap structure of the first CP layer with fitted models for CP hexamers (H0-H1) and pentamers (P1-P6).

Our results show that 6HB is suitable to direct the assembly. Its diameter matches with the central channel of empty tubes [17, 18, 37] and it simultaneously restricts the growth of long fibres. Subsequently, the effect of the origami shape on the protein layer formation was investigated. The decrease in electrophoretic mobility confirms electrostatic interactions between the origami and CPs for all DNA origami structures tested (Fig. 4a for 24HB (top) and 60HB (bottom), Supplementary Note S5, Fig. S6 for 13HR). Interestingly, the shift upon complexation for 60HB shows two distinct states while the tubular origami structures show a smoother transition. It is notable that the shift occurs at similar ratios, independent of the origami structure, being in line with a ”magic ratio” reported for the encapsulation of ssRNA [22]. The complete formation of a single layer on 24HB (Fig. 4b, middle, Supplementary Fig. S2c) is observed at *ε* = 2.5k, whereas a second layer requires *ε* = 10k (Fig. 4b, bottom, Supplementary Fig. S2d). The complexed structures are consistent in length, however, compared to 6HB, the diameter of 24HB is slightly increased (21.9±1.6 nm *vs*. 18.1±0.9 nm, s.d.). A difference in the helical symmetry is also obtained from single-particle reconstruction (Fig. 4c, Supplementary Fig. S3b, Fig. S4b). The tubes (24HB-2.5k, single layer) are again formed by hexamers arranged in a 1-start helix, however it has a turn of 48.1° and a rise of 23.5 Å. Note that, in the cross-section the highlighted 24HB origami structure (blue) is represented by two DNA layers.

**Fig. 4.**
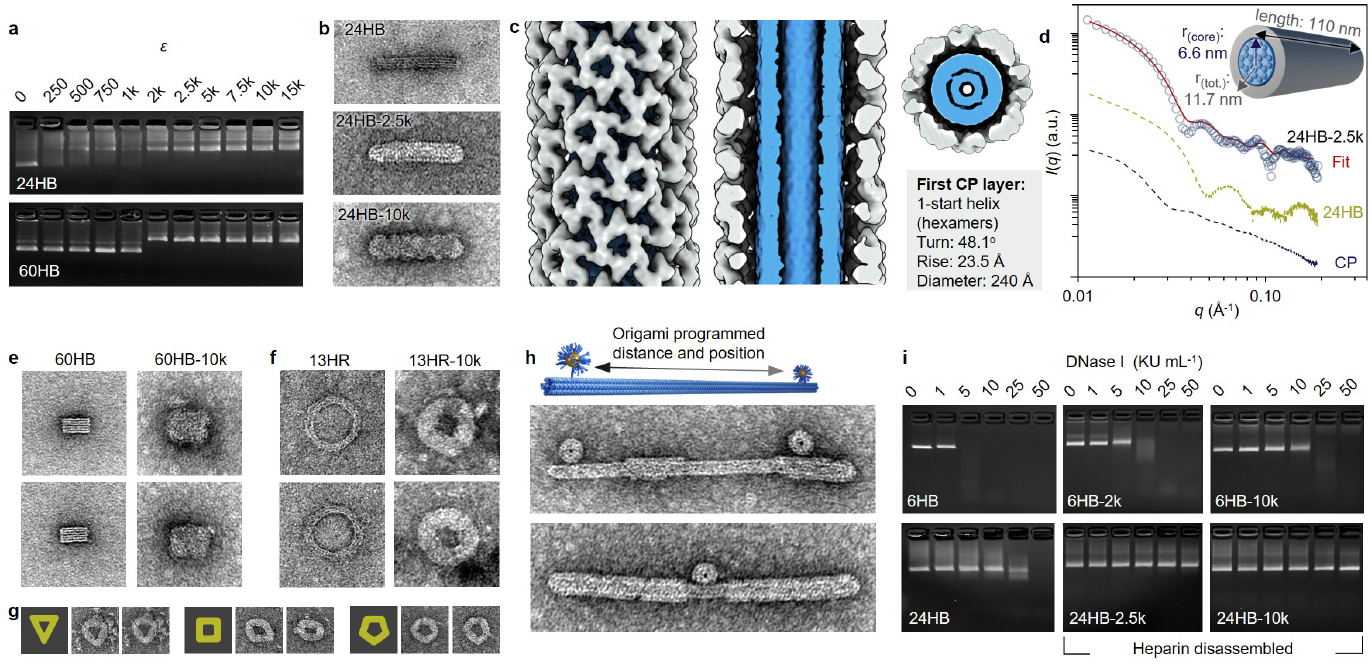
Applicability of capsid coating on structures with different thickness and shape. **a**, EMSA for 24HB (top) and 60HB (bottom) showing a decrease in the mobility with increasing *ε*. **b**, Negative-stain TEM images of plain 24HB (top) and complexed structures with *ε* = 2.5k (middle) and *ε* = 10k (bottom). The image dimensions correspond to 200 nm × 100 nm. **c**, Cryo-EM density maps for 24HB-2.5k. The origami structure is highlighted in blue. **d**, SAXS scattering curves measured for CP (blue), 24HB (green) and 24HB-2.5k samples in solution. **e**, Negative-stain TEM images (125 nm × 125 nm) of plain 60HB (left) and 60HB-10k (right). **f**, Negative-stain TEM images (125 nm × 125 nm) of 13HR structure before (left) and after (right) complexation with *ε* = 10k. **g**, Complexed 13HR (*ε* = 10k) can be classified into triangular (left), square-like (middle) and pentagonal (right) shapes. All images have dimensions of 125 nm × 125 nm. **h**, Application of protein coating on AuNP-functionalized 6HB structures. The image width corresponds to 400 nm. **i**, Stability of 6HB (top) and 24HB (bottom) upon DNase I treatment. For both structures, the digestion of the coated DNA origami (single (middle) or double protein layer (right)) is slower compared to the plain structures (left). The coating has been removed by heparin before AGE to avoid retention in the wells.

The homogeneity of the complexed structures was confirmed with smallangle X-ray scattering (SAXS, Fig. 4d). For analysis, a Debye background was added to the intensity distribution while the background in form of the complexation buffer was subtracted (Supplementary Note S6). It shows distinct curves for CPs only (dark blue) and 24HB (green), while a clear change in patterns is observed for complexed 24HB (24HB-2.5k, blue circles). A core-shell cylinder is chosen to represent the complexed structure best in a geometrical model (red). The obtained radius of 6.6 nm for the core and a shell thickness of 5.1 nm results in a total d_SAXS_ = 23.4 nm, which is in agreement with the TEM analysis and the single-particle reconstruction of 24HB-2.5k.

Successful complexation is suggested for both 60HB and 13HR from the change in electrophoretic mobility during AGE. For 60HB (Fig. 4e), due to the low aspect ratio, the formation of spherical shells rather than tubes might be expected, since the origami’s cross-section corresponds to the size of the outer diameter of tubes, which have been observed to form upon CP complexation with 6HB or 24HB. The coating of 60HB is observed to develop from the square-shaped side of the origami structure, but fully complexed structures are only observed at high ratios, with a clear trend toward smoothening of the edges (Supplementary Note S7, Fig. S8a,b). A fully developed single layer coating for the ring structure requires large *ε* (Fig. 4f; Supplementary Fig. S8c,d) although its cross-sectional size is between 6HB and 24HB, suggesting that the negative curvature is not well tolerated by the CPs. Following the tube formation along the template, clear kinks are seen, resulting in an appearance of the complexed structures predominantly as triangles, squares and pentagons (Fig. 4g). This implies that a series of short linear stretches deforming the template is more favorable for the assembly formation than a negative curvature.

Having established a system which allows precise control over the dimensions of the assembled structures (Supplementary Note S8, Fig. S9), we wanted to further exploit the properties of DNA origami. Since each strand is addressable, functional units can be precisely positioned on the origami structure. We demonstrated this with DNA coated gold nanoparticles (AuNPs) anchored on the 6HB structure (Fig. 4h, Supplementary Note S9, Fig. S10). As a result of the complexation, both the DNA origami and the AuNPs are coated, showing the formation of dumbbell-like structures with 6HB precisely controlling the distance between the spherical particles.

In addition, it is well known that the stability of DNA origami is buffer dependent, and that the origami structures are susceptible to DNase I degradation [40]. Several strategies have been developed to circumvent the degradation, including the application of different coatings [41]. Subsequently, the efficiency of the CP coating was tested for both 6HB and 24HB (Fig. 4i). Complexation with CCMV CPs is found to enhance the stability until 50 Kunitz units (KU) mL^−1^ of DNAse I (Supplementary Note S10, Fig. S11a). The complexed structures were disassembled after DNase I treatment with heparin sodium salt (Supplementary Fig. S11b), exposing the plain structures to highly active DNase I for a short time, which is, however, long enough to attack the 6HB structure.

In conclusion, we have developed a strategy to direct the assembly of virus capsids at physiological conditions in a precise and programmable manner. By employing versatile DNA origami as templates, a rigorous control over the size and shape of the formed assemblies can be achieved. The complexation is based on electrostatic interactions, which, due to the properties of the CPs, leads to the formation of one or two protein layers depending on the CP concentration used. It furthermore allows for fine-tuning the control over the assembly by altering the salt conditions or engineering of the CP-DNA interactions. Hexamers are the predominant building units of the protein layers, which were found to, dependent on their template, substantially differ in their symmetry. Although protein coatings were successfully developed onto all tested origami shapes, a preference towards tubular origami structures was observed, hence allowing even the encapsulation of DNA origami with negative curvature.

Additionally, our approach does not only enhance the stability of DNA origami against digestion by DNase I, but provides a highly functionalizable platform. The high addressability of DNA origami, which was demonstrated with AuNPs, enables precise attachment of a variety of cargo/targeting molecules by hybridization or intercalation. In combination with functionalization of the protein component, the resulting multipurpose system could be implemented in various fields ranging from materials science to biomedicine.

## Methods

### DNA origami folding and purification

The DNA origami was folded in a one-pot reaction by gradually decreasing the temperature using a Proflex 3×32-well PCR system (ThermoFisher). The scaffold strands (p7249 and p7560 variants of single-stranded M13mp18) were purchased from Tilibit Nanosystems and the staple strands from Integrated DNA Technologies. For 6HB [42], 60HB [43], and 13HR [44] the p7249 scaffold was used. 24HB [45, 46], for one, was folded from p7560 (Supplementary Note S1).

Briefly, all structures with exception of 13HR were folded using final concentrations of 20 nM scaffold and 200 nM of each staple strand in a buffered environment (”folding buffer”, FOB). The FOB contains 1× Tris-acetate-ethylenediaminetetraacetic acid (EDTA) (1× TAE pH 8.4, 40 mM Tris, 20 mM acetic acid and 1 mM EDTA) which was supplemented with varying salt concentrations depending on the DNA origami structure: 12.5 mM MgCl_2_ for 6HB, 17.5 mM MgCl_2_ for 24HB, and 20 mM MgCl_2_ with 5 mM NaCl for 60HB. The following thermal annealing ramp was used for 24HB and 60HB folding: from 65 °C to 59 °C at a rate of −4 °C h^−1^, and from 59 °C to 40 °C at −0.33 °C h^−1^. For 6HB, a different annealing protocol was used: from 90 °C to 70 °C at a rate of −1.5 °C min^−1^, from 70 °C to 60 °C at −0.75 °C min^−1^, and from 60 °C to 27 °C at −3 °C h^−1^.

These above-mentioned structures were purified from excess staple strands using poly(ethylene glycol) (PEG) precipitation [47]. After dilution in 1 × FOB to a concentration of approx. 5 nM, the final volume was mixed in a 1:1 ratio with PEG precipitation buffer (1× TAE, 15 % (w/v) PEG8000, 505 mM NaCl) and centrifuged for 30 min at 14,000 *g*. The supernatant was removed, and the pelleted DNA origami resuspended in 1 × FOB and incubated overnight at 30 °C, 600 rpm on an Eppendorf ThermoMixer C.

13HR was folded with final scaffold and staple concentrations of 10 nM and 50 nM, respectively, in 1× Tris-EDTA (0× TE pH 7.6, 10 mM Tris, 1 mM EDTA) buffer containing 10 mM MgCl_2_. After 15 min incubation at 80 °C, the mixture was cooled from 79 °C to 71 °C at a rate of −1 °C min^−1^, from 70 °C to 66 °C at −0.2 °C min^−1^, from 65 °C to 60 °C at −2 °C h^−1^, from 59 °C to 37 °C at −1 °C h^−1^, from 36 °C to 30 °C at −4 °C h^−1^ and from 29 °C to 20 °C at −0.2 °C min^−1^. Due to dimer formation and aggregation of 13HR, this structure was purified from an agarose gel. To this end, the samples were loaded on a 1 % (w/v) agarose gel (0.5 × Tris-borate-EDTA (TBE) buffer, pH 8.3, containing 44.5 mM Trizma base, 44.5 mM boric acid, 1 mM EDTA supplemented with 11 mM MgCl_2_) and run for 2.25 h at 80 V. For visualization, ethidium bromide (final concentration of 0.46 μg mL^−1^) was used and the target band was cut out under ultraviolet (UV) light. The gel was cut in small pieces which were added into ”freeze’n’squeeze” cups (BioRad) and frozen for 5 min. Subsequently, the DNA origami was recovered after centrifugation at 16,000 *g* and 10 °C for 10 min.

The origami concentration was estimated according to Lambert-Beer’s law from the absorbance measured at 260 nm (BioTek Eon Microplate Spectrophotometer, 2 μL sample volume, Take3 plate). The extinction coefficient of DNA origami structures is dependent on the number of hybridized and non-hybridized nucleotides [48] and was estimated to be 0.98 × 10^8^ M^−1^ cm^−1^ for 6HB, 1.076 × 10^8^ M^−1^ cm^−1^ for 24HB, 0.91 × 10^8^ M^−1^ cm^−1^ for 60HB, and 1.3 × 10^8^ M^−1^ cm^−1^ for 13HR.

### Buffer exchange for DNA origami

The purified DNA origami structures were transferred into 6.5 mM 4-(2-hydroxyethyl)-1-piperazineethanesulfonic acid (HEPES) buffer supplemented with 2 mM NaOH (HEPES-NaOH, pH 6.5) before complexation. The buffer exchange was performed by spin-filtration [49] using 100 kDa molecular weight cut-off (MWCO) centrifugal filters (Amicon), which were washed before use by centrifuging with 400 μL of HEPES-NaOH buffer for 5 min at 14,000 *g*. Subsequently, equal volumes of DNA origami solution and HEPES-NaOH buffer were added into the filter device and the centrifugation was continued for 10 min at 6,000 *g*. HEPES-NaOH was then added in the quantity of 2.09× the initial volume of origami solution, and the centrifugation step was repeated. The sample was collected by inverting the filter and centrifuging for 2.5 min at 1,000 *g*.

### Isolation of CCMV capsid proteins

The CPs were isolated from intact CCMV, similar to a protocol by Mikkilä *et al*. [50]. First, the virus was dialysed overnight against 50 mM Tris-HCl, 500 mM CaCl_2_ buffer, pH 7.5 supplemented with 1 mM dithiothreitol (DTT) using Slize-A-Lyzer Mini Dialysis cups (3.5 kDa MWCO, Thermo Scientific). The RNA was pelleted in a centrifugation step at 4 °C using 21,100 *g* for 6 h and the recovered supernatant was dialysed overnight against ”clean buffer” which contains 50 mM Tris-HCl, 150 mM NaCl at pH 7.5 supplemented with 1 mM DTT. The concentration of the proteins was determined based on their absorbance at 280 nm (extinction coefficient of 23,590 M^−1^ cm^−1^) using a BioTek Eon microplate spectrophotometer (2 μL sample, Take3 plate).

### Agarose gel electrophoresis

Agarose gel electrophoresis (AGE) was used to study the binding interaction between the proteins and the origami structures by monitoring the shift in electrophoretic mobility. Furthermore, the intactness of the origami structures after folding and purification, and during DNase I digestion was analyzed by gel electrophoresis. To this end samples (volumes ranging from 10-32 μL) supplemented with 6x gel loading dye (40% sucrose w/o dye for samples from digestion studies) were run in a 2 % (w/v) agarose gel (1× TAE buffer, 11 mM MgCl_2_) for 45 min at 90 V in 1× TAE buffer supplemented with 11 mM MgCl_2_. For staining, ethidium bromide at a final concentration of 0.46 μg mL^−1^ was used and the DNA was visualized under UV light using a GelDoc XR+ system (Bio-Rad).

### Complexation of DNA origami and capsid proteins

The complexation between CPs and DNA origami was performed at a final origami concentration of 4 nM (10 μL samples). The origami was added in a 1:1 volume ratio to the protein solution which has been diluted in the ”clean buffer”. Depending on the required protein excess, *ε*, which describes the molar ratio between CP to DNA origami, protein solutions ranging from 0–60 μM (corresponding to *ε* = 0–15k) have been prepared. The NaCl concentration was adjusted to 150 mM, resulting in a complexation buffer containing 3.25 mM HEPES-NaOH, 25 mM Tris-HCl, 150 mM NaCl and 0.5 mM DTT. The complexation was performed at 4 °C for at least 1 h and subsequently analysed using AGE and TEM.

### DNase I digestion assays

To study the protection effect of the CP coating against degradation of the origami structures by DNase I, 2 μL of DNase I stock (ranging from 0–500 KU mL^−1^ were added to 16 μL of the sample. Additionally, CaCl_2_ and MgCl_2_ concentrations were adjusted, resulting in a final reaction volume of 20 μL containing 3.2 nM DNA origami, 2.6 mM HEPES-NaOH, 20 mM Tris-HCl, 120 mM NaCl, 0.4 mM DTT, 1 mM CaCl_2_ and 5 mM MgCl_2_. The samples are incubated at 37 °C for 15 min (6HB) and 60 min (24HB). Before the outcome is analysed using AGE, samples complexed with CPs were disassembled using heparin sodium salt as a competitive binding agent (final concentrations of 1.5 μM for 6HB-2k and 24HB-2.5k and 82 μM for 6HB-10k and 24HB-10k, Supplementary Note S10).

### Transmission electron microscopy

Plain origami samples (4 nM) were prepared by incubation of a 3 μL droplet for 3 min on a plasma cleaned (20 s oxygen plasma flash, Gatan solarus) Formvar carbon-coated copper grid (FCF400Cu, Electron Microscopy Sciences) which was subsequently blotted against filter paper and negative stained, similar to the protocol described by Castro *et al*. [51]. For complexed samples (4 nM), a 3 μL droplet was deposited for 1.5 min. After blotting against filter paper, the grid was immersed into a 10 μL droplet of complexation buffer (3.25 mM HEPES-NaOH, 25 mM Tris-HCl, 150 mM NaCl, 0.5 mM DTT) for 5 s. Negative staining was performed in aqueous 2 % (w/v) uranyl formate solution (supplemented with 25 mM NaOH for pH adjustment) by immersing the grid in a 5 μL droplet which was immediately blotted away. It was followed by an immersion into a 20 μL droplet which was incubated on the grid for 45 s. After the final blotting step, the samples were left for drying for at least 20 min before imaging was performed on a FEI Tecnai 12 Bio-Twin microscope at an acceleration voltage of 120 V.

### Cryo-electron microscopy

The samples for cryo-EM were prepared using a vitrification apparatus (Vitrobot, Thermo Fisher Scientific). The origami concentrations in the complexed samples were 90 nM for 6HB-2k, 84 nM for 24HB-2.5k, 18 nM for 6HB-10k, and 21 nM for 24HB-10k, resulting in a total CP concentration of 180 μM and 210 μM for complexed 6HB and 24HB samples, respectively. A 3 μL aliquot of the complexed origami sample was deposited on a plasma cleaned (50 s, Harrick Plasma PDC-002-EC instrument) holey carbon-coated grid (copper 200 mesh R1.2/1.3, Quantifoil). After a 1 min incubation, excess liquid was blotted for 10 s at 100 % relative humidity and 6 °C, followed by plunging the grid into liquid ethane. The grids were stored in liquid nitrogen. Data were collected on a Talos Arctica transmission electron microscope (Thermo Fisher Scientific) at liquid nitrogen temperature operated at 200 kV, using a Falcon III direct electron detector (Thermo Fisher Scientific). A magnification of 150,000× was used, resulting in a calibrated pixel size of 0.96 Å. The data collection parameters are listed in Table S1 (Supplementary Note S11).

### Single-particle reconstruction

Cryo-EM data were processed using CryoSPARC 3.3.2 (Structura Biotechnology) unless stated otherwise. Contrast transfer function parameters were estimated using CTFFIND4 [52]. Segments along filaments were defined using the Filament Tracer function. Helical symmetry parameters were estimated initially from 2D class averages using Python-based Helix Indexer [53]. The structure and helical symmetry parameters were refined using Helix Refine function and non-uniform refinement on motion corrected helix segments. To determine the helical symmetry parameters of the 6HB-10k outer layer, a second 2D classification run was performed after subtracting the contribution of the inner layer using the Particle Subtraction function. The Helix Refine was ran on the subset of particles that showed a clear second layer, using the determined symmetry parameters as initial estimates. Reconstructions were sharpened by applying an ad hoc B-factor of –300 Å^2^. The reconstructions were averaged in real space by imposing the helical symmetry parameters on the central, most ordered part of the map (50 % of the volume) in Bsoft [54]. For modelling the structure of the capsomer, CP monomer (PDB:1cwp) was fitted in the 6HB-2k reconstruction in the six positions of the hexamer as rigid bodies in UCSF ChimeraX 1.3 [55]. The atomic model was refined against the density using ISOLDE 1.3 [56] and Phenix 1.19 [57]. To create atomic representations of the filaments, symmetry copies of the hexamer were created in ChimeraX. To visualize the placement of CP hexamers and pentamers in the cap, the caps of the 6HB-2k filament were manually picked in the micrographs. The cap structure was refined using the Helix Refine function omitting symmetrization, as this allowed limiting the tilt angle of the caps close to side-views. Reconstruction of the cap was filtered to its local resolution using Local Filter. The hexamer atomic model and previously determined pentamer structure (extracted from PDB:1cwp after applying icosahedral symmetry) were fitted as rigid bodies in ChimeraX 1.3. Data processing parameters are given in Table S1. Model refinement and validation parameters are shown in Table S2.

### Small-angle X-ray scattering

The samples for SAXS were prepared at an origami concentration of 180 nM (corresponding to a disassembled CP concentration of 450 μM) and sealed within a 1.5 mm diameter glass capillary. The measurements were performed using a Xenocs Xeuss 3.0 C device equipped with a GeniX 3D Cu microfocus source (wavelength λ = 1.542 Å) and EIGER2 R 1M hybrid pixel detector at a sample-detector distance of 1100 mm. Data acquisition was performed for 3× 3 h per sample. To obtain the one-dimensional SAXS data, the 2D scattering data was azimuthally averaged. The magnitude of the scattering vector *q* is given by *q* = 4πsinθ/λ with 2θ being the scattering angle. Data treatment included averaging of the triplicate 2D data of each sample, background subtraction from the complexation buffer (3.25 mM HEPES-NaOH, 25 mM Tris-HCl, 150 mM NaCl, 0.5 mM DTT) and a form factor was fitted to a cylinder (24HB), spheres (T = 3 icosahedral CPs assemblies) and a core-shell cylinder (24HB-2.5k) using SasView software. A Debye Anderson Brumberger model was added to account for the background.

## Supplementary Information

A collection of Supplementary Notes (Notes 1–11) including Supplementary Figures (S1–11) and Supplementary Tables (S1–2) is found in the accompanying .pdf file.

## Acknowledgments

The authors acknowledge financial support from the European Research Council (ERC) and ERA Chair MATTER under European Union’s Horizon 2020 research and innovation programme (grant agreements no. 101002258 and no. 856705), Emil Aaltonen Foundation, Sigrid Jusélius Foundation, Academy of Finland (grant no. 341057) and Jane and Aatos Erkko Foundation. Kha M. Nguyen and Anton Kuzyk are acknowledged for providing the staple strands for the 13HR sample. We thank Pasi Laurinmäki and Benita Löflund for technical assistance in cryo-EM. The facilities and expertise of the HiLIFE cryo-EM unit at the University of Helsinki, a member of Instruct-ERIC Centre Finland, FINStruct, and Biocenter Finland are gratefully acknowledged. The authors wish to acknowledge CSC - IT Center of Science, Finland, for computational resources. This work was carried out under the Academy of Finland Centers of Excellence Program (2022-2029) in Life-Inspired Hybrid Materials (LIBER), project number (346110). We thank for the provision of facilities and technical support by Aalto University Bioeconomy Facilities, OtaNanoNanomicroscopy Center (Aalto-NMC) and Micronova Nanofabrication Center.

## References

[1] Golub, E., Subramanian, R.H., Esselborn, J., Alberstein, R.G., Bailey, J. B., Chiong, J.A., Yan, X., Booth, T., Baker, T.S., Tezcan, F.A.: Constructing protein polyhedra via orthogonal chemical interactions. Nature 578(7793), 172–176 (2020). https://doi.org/10.1038/s41586-019-1928-2

[2] Malay, A.D., Miyazaki, N., Biela, A., Chakraborti, S., Majsterkiewicz, K., Stupka, I., Kaplan, C.S., Kowalczyk, A., Piette, B.M.A.G., Hochberg, G.K.A., Wu, D., Wrobel, T.P., Fineberg, A., Kushwah, M.S., Kelemen, M., Vavpetič, P., Pelicon, P., Kukura, P., Benesch, J.L.P., Iwasaki, K., Heddle, J.G.: An ultra-stable gold-coordinated protein cage displaying reversible assembly. Nature 559(7756), 438–442 (2019). https://doi.org/10.1038/s41586-019-1185-4

[3] Hsia, Y., Bale, J.B., Gonen, S., Shi, D., Sheffler, W., Fong, K.K., Nattermann, U., Xu, C., Huang, P.-S., Ravichandran, R., Yi, S., Davis, T.N., Gonen, T., King, N.P., Baker, D.: Design of a hyperstable 60-subunit protein icosahedron. Nature 535(7610), 136–139 (2016). https://doi.org/10.1038/nature18010

[4] Bale, J.B., Gonen, S., Liu, X., Sheffler, W., Ellis, D., Thomas, C., Cascio, D., Yeates, T.O., Gonen, T., King, N.P., Baker, D.: Accurate design of megadalton-scale two-component icosahedral protein complexes. Science 353(6297), 389–394 (2016). https://doi.org/10.1126/science.aaf8818

[5] Marcandalli, J., Fiala, B., Ols, S., Perotti, M., de van der Schueren, W., Snijder, J., Hodge, E., Benhaim, M., Ravichandran, R., Carter, L., Sheffler, W., Brunner, L., Lawrenz, M., Dubois, P., Lanzavecchia, A., Sal-lusto, F., Lee, K.K., Veesler, D., Correnti, C.E., Stewart, L.J., Baker, D., Lorè, K., Perez, L., King, N.P.: Induction of potent neutralizing antibody response by a designed protein nanoparticle vaccine for respiratory syncytial virus. Cell 176(6), 1420–1431 (2019). https://doi.org/10.1016/j.cell.2019.01.046

[6] Hartman, E.C., Jakobson, C.M., Favor, A.H., Lobba, M.J., Álvarez-Benedicto, E., Francis, M.B., Tullman-Ercek, D.: Quantitative characterization of all single amino acid variants of a viral capsid-based drug delivery vehicle. Nat. Commun. 9, 1385 (2018). https://doi.org/10.1038/s41467-018-03783-y

[7] Douglas, T., Young, M.: Host-guest encapsulation of materials by assembled virus protein cages. Nature 393(6681), 152–155 (1998). https://doi.org/10.1038/30211

[8] Comellas-Aragonès, M., Engelkamp, H., Claessen, V.I., Sommerdijk, N.A.J.M., Rowan, A.E., Christianen, P.C.M., Maan, J.C., Verduin, B.J.M., Cornelissen, J.J.L.M., Nolte, R.J.M.: A virus-based single-enzyme nanoreactor. Nat. Nanotechnol. 2(10), 635–639 (2007). https://doi.org/10.1038/nnano.2007.299

[9] Mao, C., Solis, D.J., Reiss, B.D., Kottmann, S.T., Sweeney, R.Y., Hayhurst, A., Georgiou, G., Iverson, B., Belcher, A.M.: Virus-based toolkit for the directed synthesis of magnetic and semiconducting nanowires. Science 303(5655), 213–217 (2004). https://doi.org/10.1126/science.1092740

[10] Lizotte, P.H., Wen, A.M., Sheen, M.R., Fields, J., Rojanasopondist, P., Steinmetz, N.F., Fiering, S.: *In situ* vaccination with cowpea mosaic virus nanoparticles suppresses metastatic cancer. Nat. Nanotechnol. 11(3), 295–303 (2015). https://doi.org/10.1038/nnano.2015.292

[11] Ehrlich, L.S., Agresta, B.E., Carter, C.A.: Assembly of recombinant human immunodeficiency virus type 1 capsid protein in vitro. J. Virol. 66(8), 4874–4883 (1992). https://doi.org/10.1128/jvi.66.8.4874-4883.1992

[12] Li, S., Hill, C.P., Sundquist, W.I., Finch, J.T.: Image reconstruction of helical assemblies of the HIV-1CA protein. Nature 407(6802), 409–413 (2000). https://doi.org/10.1038/35030177

[13] Zheng, C., Lázaro, G.R., Tsvetkova, I.B., Hagan, M.F., Dragnea, B.: Defects and chirality in the nanoparticle-directed assembly of spherocylindrical shells of virus coat proteins. ACS Nano 12(6), 5323–5332 (2018). https://doi.org/10.1021/acsnano.8b00069

[14] Dhason, M.S., Wang, J.C.-Y., Hagan, M.F., Zlotnick, A.: Differential assembly of Hepatitis B Virus core protein on single-and double-stranded nucleic acid suggest that dsDNA-filled core is spring-loaded. Virology 430(1), 20–29 (2012). https://doi.org/10.1016/j.virol.2012.04.012

[15] Biela, A.P., Naskalska, A., Fatehi, F., Twarock, R., Heddle, J.G.: Programmable polymorphism of a virus-like particle. Commun. Mater. 3, 7 (2022). https://doi.org/10.1038/s43246-022-00229-3

[16] Sun, J., DuFort, C., Daniel, M.-C., Murali, A., Chen, C., Gopinath, K., Stein, B., De, M., Rotello, V.M., Holzenburg, A., Kao, C.C., Dragnea, B.: Core-controlled polymorphism in virus-like particles. Proc Natl Acad Sci U S A. 104(4), 1354–1359 (2007). https://doi.org/10.1073/pnas.06105421

[17] Speir, J.A., Munshi, S., Wang, G., Baker, T.S., Johnson, J.E.: Structures of the native and swollen forms of cowpea chlorotic mottle virus determined by X-ray crystallography and cryo-electron microscopy. Structure 3(1), 63–78 (1995). https://doi.org/10.1016/S0969-2126(01)00135-6

[18] Bancroft, J.B., Hills, G.J., Markham, R.: A study of the self-assembly process in a small spherical virus. Formation of organized structures from protein subunits in vitro. Virology 31(2), 354–379 (1967). https://doi.org/10.1016/0042-6822(67)90180-8

[19] Bancroft, J.B., Bracker, C.E., Wagner, G.W.: Structures derived from cowpea chlorotic mottle and brome mosaic virus protein. Virology 38(2), 324–335 (1969). https://doi.org/10.1016/0042-6822(69)90374-2

[20] Adolph, K.W., Butler, P.J.G.: Studies on the assembly of a spherical plant virus. I. States of aggregation of the isolated protein. J. Mol. Biol. 88(2), 327–341 (1974). https://doi.org/10.1016/0022-2836(74)90485-9

[21] Lavelle, L., Gingery, M., Phillips, M., Gelbart, W.M., Knobler, C.M., Cadena-Nava, R.D., Vega-Acosta, J.R., Pinedo-Torres, L.A., Ruiz-Garcia, J.: Phase diagram of self-assembled viral capsid protein polymorphs. J. Phys. Chem. B 113(12), 3813–3819 (2009). https://doi.org/10.1021/jp8079765

[22] Comas-Garcia, M., Garmann, R.F., Singaram, S.W., Ben-Shaul, A., Knobler, C.M., Gelbart, W.M.: Characterization of viral capsid protein self-assembly around short single-stranded RNA. J. Phys. Chem. B 118(27), 7510–7519 (2014). https://doi.org/10.1021/jp503050z

[23] Chang, C.B., Knobler, C.M., Gelbart, W.M., Mason, T.G.: Curvature dependence of viral protein structures on encapsidated nanoemulsion droplets. ACS Nano 2(2), 281–286 (2008). https://doi.org/10.1021/nn700385z

[24] Aniagyei, S.E., Kennedy, C.J., Stein, B., Willits, D.A., Douglas, T., Young, M.J., De, M., Rotello, V.M., Srisathiyanarayanan, D., Kao, C.C., Dragnea, B.: Synergistic effects of mutations and nanoparticle templating in the self-assembly of cowpea chlorotic mottle virus capsids. Nano Lett. 9(1), 393–398 (2009). https://doi.org/10.1021/nl8032476

[25] Rothemund, P.: Folding DNA to create nanoscale shapes and patterns. Nature 440(7082), 297–302 (2006). https://doi.org/10.1038/nature04586

[26] Dietz, H., Douglas, S.M., Shih, W.M.: Folding DNA into twisted and curved nanoscale shapes. Science 325(5941), 725–730 (2009). https://doi.org/10.1126/science.1174251

[27] Douglas, S.M., Dietz, H., Liedl, T., Högberg, B., Graf, F., Shih, W.M.: Self-assembly of DNA into nanoscale three-dimensional shapes. Nature 459(7245), 414–418 (2009). https://doi.org/10.1038/nature08016

[28] Dey, S., Fan, C., Gothelf, K.V., Li, J., Lin, C., Liu, L., Liu, N., Nijenhuis, M.A., Saccà, B., Simmel, F.C., Yan, H., Zhan, P.: DNA origami. Nat. Rev. Methods Primers 1(1), 13 (2021). https://doi.org/10.1038/s43586-020-00009-8

[29] Villalta, A., Schmitt, A., Estrozi, L.F., Quemin, E.R.J., Alempic, J.-M., Lartigue, A., Pražák, V., Belmudes, L., Vasishtan, D., Colmant, A.M.G., Honoré, F.A., Couté, Y., Grünewald, K., Abergel, C.: The giant mimivirus 1.2 Mb genome is elegantly organized into a 30 nm diameter helical protein shield. eLife 11, 77607 (2022). https://doi.org/10.7554/eLife.77607

[30] Johnson, J.M., Willits, D.A., Young, M.J., Zlotnick, A.: Interaction with capsid protein alters RNA structure and the pathway for *in vitro* assembly of cowpea chlorotic mottle virus. J. Mol. Biol. 335(2), 455–464 (2004). https://doi.org/10.1016/j.jmb.2003.10.059

[31] Zlotnick, A., Aldrich, R., Johnson, J.M., Ceres, P., Young, M.J.: Mechanism of capsid assembly for an icosahedral plant virus. Virology 277(2), 450–456 (2000). https://doi.org/10.1006/viro.2000.0619

[32] Hespenheide, B.M., Jacobs, D.J., Thorpe, M.F.: Structural rigidity in the capsid assembly of cowpea chlorotic mottle virus. J. Phys. Condens. Matter 16(44), 5055 (2004). https://doi.org/10.1088/0953-8984/16/44/003

[33] Endres, D., Zlotnick, A.: Model-based analysis of assembly kinetics for virus capsids or other spherical polymers. Biophys. J. 83(2), 1217–1230 (2002). https://doi.org/10.1016/S0006-3495(02)75245-4

[34] McPherson, A.: Micelle formation and crystallization as paradigms for virus assembly. BioEssays 27(4), 447–458 (2005). https://doi.org/10.1002/bies.20196

[35] Perlmutter, J.D., Perkett, M.R., Hagan, M.F.: Pathways for virus assembly around nucleic acids. J. Mol. Biol. 426(18), 3148–3165 (2014). https://doi.org/10.1016/j.jmb.2014.07.004

[36] Lázaro, G.R., Dragnea, B., Hagan, M.F.: Self-assembly of convex particles on spherocylindrical surfaces. Soft Matter 14(28), 5728–5740 (2018). https://doi.org/10.1039/C8SM00129D

[37] Mukherjee, S., Pfeifer, C.M., Johnson, J.M., Liu, J., Zlotnick, A.: Redirecting the coat protein of a spherical virus to assemble into tubular nanostructures. J. Am. Chem. Soc. 128(8), 2538–2539 (2006). https://doi.org/10.1021/ja056656f

[38] de Ruiter, M.V., van der Hee, R.M., Driessen, A.J.M., Keurhorst, E.D., Hamid, M., Cornelissen, J.J.L.M.: Polymorphic assembly of virus-capsid proteins around DNA and the cellular uptake of the resulting particles. J. Control Release 307, 342–354 (2019). https://doi.org/10.1016/j.jconrel.2019.06.019

[39] de la Escosura, A., Janssen, P.G.A., Schenning, A.P.H.J., Nolte, R.J.M., Cornelissen, J.J.L.M.: Encapsulation of DNA-templated chromophore assemblies within virus protein nanotubes. Angew. Chem. Int. Ed 49(31), 5335–5338 (2010). https://doi.org/10.1002/anie.201001702

[40] Ramakrishnan, S., Ijäs, H., Linko, V., Keller, A.: Structural stability of DNA origami nanostructures under application-specific conditions. Comput. Struct. Biotechnol. J. 16, 342–349 (2018). https://doi.org/10.1016/j.csbj.2018.09.002

[41] Bila, H., Kurisinkal, E.E., Bastings, M.M.C.: Engineering a stable future for DNA-origami as a biomaterial. Biomater. Sci. 7(2), 532–541 (2019). https://doi.org/10.1039/C8BM01249K

[42] Bui, H., Onodera, C., Kidwell, C., Tan, Y., Graugnard, E., Kuang, W., Lee, J., Knowlton, W.B., Yurke, B., Hughes, W.L.: Programmable periodicity of quantum dot arrays with DNA origami nanotubes. Nano Lett. 10(9), 3367–3372 (2010). https://doi.org/10.1021/nl101079u

[43] Linko, V., Shen, B., Tapio, K., Toppari, J.J., Kostiainen, M.A., Tuukkanen, S.: One-step large-scale deposition of salt-free DNA origami nanostructures. Sci. Rep. 5, 15634 (2015). https://doi.org/10.1038/srep15634

[44] Nguyen, M.-K., Nguyen, V.H., Natarajan, A.K., Huang, Y., Ryssy, J., Shen, B., Kuzyk, A.: Ultrathin silica coating of DNA origami nanostructures. Chem. Mater. 32(15), 6657–6665 (2020). https://doi.org/10.1021/acs.chemmater.0c02111

[45] Ijäs, H., Shen, B., Heuer-Jungemann, A., Keller, A., Kostiainen, M.A., Liedl, T., Ihalainen, J.A., Linko, V.: Unraveling the interaction between doxorubicin and DNA origami nanostructures for customizable chemotherapeutic drug release. Nucleic Acids Res. 49(6), 3048–3062 (2021). https://doi.org/10.1093/nar/gkab097

[46] Seitz, I., Ijäs, H., Linko, V., Kostiainen, M.A.: Optically responsive protein coating of DNA origami for triggered antigen targeting. ACS Appl. Mater. Interfaces 14(34), 38515–38524 (2022). https://doi.org/10.1021/acsami.2c10058

[47] Stahl, E., Martin, T.G., Praetorius, F., Dietz, H.: Facile and scalable preparation of pure and dense DNA origami solutions. Angew. Chem. Int. Ed. 53(47), 12735–12740 (2014). https://doi.org/10.1002/anie.201405991

[48] Hung, A.M., Micheel, C.M., Bozano, L.D., Osterbur, L.W., Wallraff, G.M., Cha, J.N.: Large-area spatially ordered arrays of gold nanoparticles directed by lithographically confined DNA origami. Nat. Nanotechnol. 5(2), 121–126 (2010). https://doi.org/10.1038/nnano.2012.237

[49] Julin, S., Nonappa, Shen, B., Linko, V., Kostiainen, M.A.: DNA-origami-templated growth of multilamellar lipid assemblies. Angew. Chem. Int. Ed. 60(2), 827–833 (2021). https://doi.org/10.1002/anie.202006044

[50] Mikkilä, J., Eskelinen, A.-P., Niemelä, E.H., Linki, V., Frilander, M.J., Täormaä, P., Kostiainen, M.A.: Virus-encapsulated DNA origami nanostructures for cellular delivery. Nano Lett. 14(4), 2196–2200 (2014). https://doi.org/10.1021/nl500677j

[51] Castro, C.E., Kilchherr, F., Kim, D.-N., Shiao, E.L., Wauer, T., Wortmann, P., Bathe, M., Dietz, H.: A primer to scaffolded DNA origami. Nat. Methods 8(3), 221–229 (2011). https://doi.org/10.1038/nmeth.1570

[52] Rohou, A., Grigorieff, N.: CTFFIND4: Fast and accurate defocus estimation from electron micrographs. J. Struct. Biol. 192(2), 216–221 (2015). https://doi.org/10.1016/j.jsb.2015.08.008

[53] Zhang, X.: Python-based Helix Indexer: A graphical user interface program for finding symmetry of helical assembly through Fourier–Bessel indexing of electron microscopic data. Protein Sci. 31(1), 107–117 (2022). https://doi.org/10.1002/pro.4186

[54] Heymann, J.B.: Guidelines for using Bsoft for high resolution reconstruction and validation of biomolecular structures from electron micrographs. Protein Sci. 27(1), 159–171 (2018). https://doi.org/10.1002/pro.3293

[55] Pettersen, E.F., Goddard, T.D., Huang, C.C., Meng, E.C., Couch, G.S., Croll, T.I., Morris, J.H., Ferrin, T.E.: USFC ChimeraX: Structure visualization for researchers, educators, and developers. Protein Sci. 30(1), 70–82 (2021). https://doi.org/10.1002/pro.3943

[56] Croll, T.I.: *ISOLDE*: a physically realistic environment for model building into low-resolution electron-density maps. Acta Crystallogr. D Struct. Biol. 74(6), 519–530 (2018). https://doi.org/10.1107/S2059798318002425

[57] Afonine, P.V., Poon, B.K., Read, R.J., Sobolev, O.V., Terwilliger, T.C., Urzhumtsev, A., Adams, P.D.: Real-space refinement in *PHENIX* for cryo-EM and crystallography. Acta Crystallogr. D Struct. Biol. 74(6), 531–544 (2018). https://doi.org/10.1107/S2059798318006551

