## Supplementary Information for "DNA origami directed virus capsid polymorphism"

### Supplementary Information for
### DNA origami directed virus capsid polymorphism

**Iris Seitz[1], Esa-Pekka Kumpula[2], Eduardo Anaya-Plaza[1], Jeroen J. L. M. Cornelissen[3], Veikko Linko[1,4,5], Juha T. Huiskonen[2*] & Mauri A. Kostiainen[1,4*]**

[1]Biohybrid Materials, Department of Bioproducts and Biosystems, Aalto University, P.O. Box 16100, 00076 Aalto, Finland

[2]Institute of Biotechnology, Helsinki Institute of Life Science HiLIFE, University of Helsinki, Helsinki, Finland

[3]Department of Molecules and Materials, MESA+ Institute for Nanotechnology, University of Twente, 7500 AE Enschede, The Netherlands

[4]LIBER Center of Excellence, Aalto University, P.O. Box 16100, 00076 Aalto, Finland

[5]Institute of Technology, University of Tartu, Nooruse 1, 50411 Tartu, Estonia

#### Contents


**Page**

Note S1: Folding of DNA origami structures
(Supplementary Figure 1) . . . . . S3
Note S2: Supplementary cryo-EM micrographs
(Supplementary Figure 2) . . . . . S4
Note S3: Single-particle reconstruction
(Supplementary Figures 3–4) . . . . . S5
Note S4: Reconstruction of the cap structure
(Supplementary Figure 5) . . . . . S7
Note S5: Coating of 13HR structure
(Supplementary Figure 6) . . . . . S8
Note S6: SAXS analysis
(Supplementary Figure 7) . . . . . S9
Note S7: Negative-stain TEM images
(Supplementary Figure 8) . . . . . S10
Note S8: Size distribution of coated structures
(Supplementary Figure 9) . . . . . S11
Note S9: Functionalization of 6HB
(Supplementary Figure 10) . . . . . S11
Note S10: DNase I digestion studies
(Supplementary Figure 11) . . . . . S13
Note S11: Collection of parameters used in cryo-EM and single-particle reconstruction
(Supplementary Tables 1–2) . . . . . S13
Supplementary Information references . . . . . S14

#### Note S1: Folding of DNA origami structures

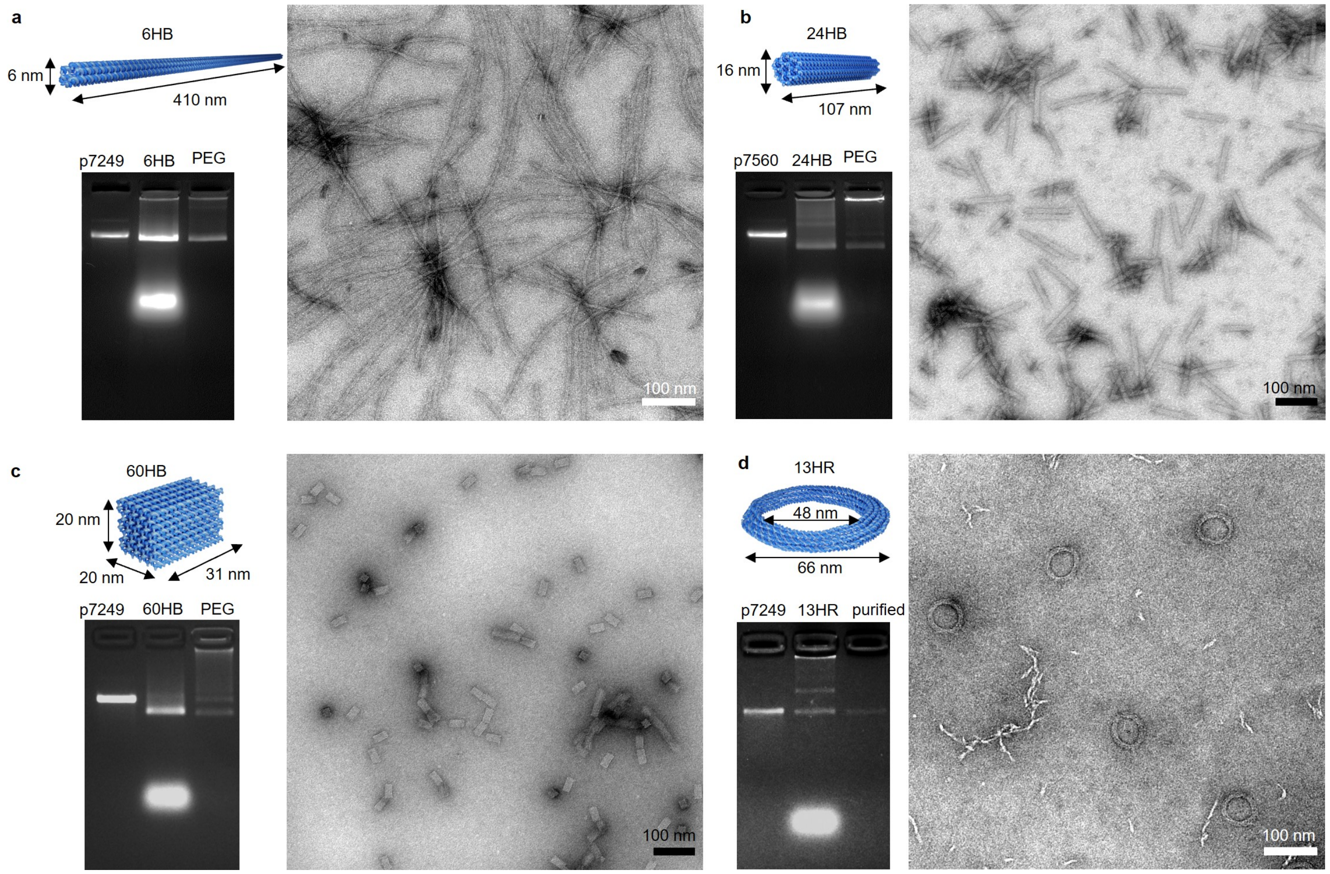


**Fig. S1** Characterization of the DNA origami structures, **a**, 6HB, **b**, 24HB, **c**, 60HB and **d**, 13HR. For each structure, a schematic showing the dimensions (top left) and the corresponding TEM micrograph (right) are shown. Agarose gel electrophoresis (AGE, bottom right) shows the folded structures before (lane 2) and after purification (lane 3) in comparison to the scaffold (lane 1).

#### Note S2: Supplementary cryo-EM micrographs

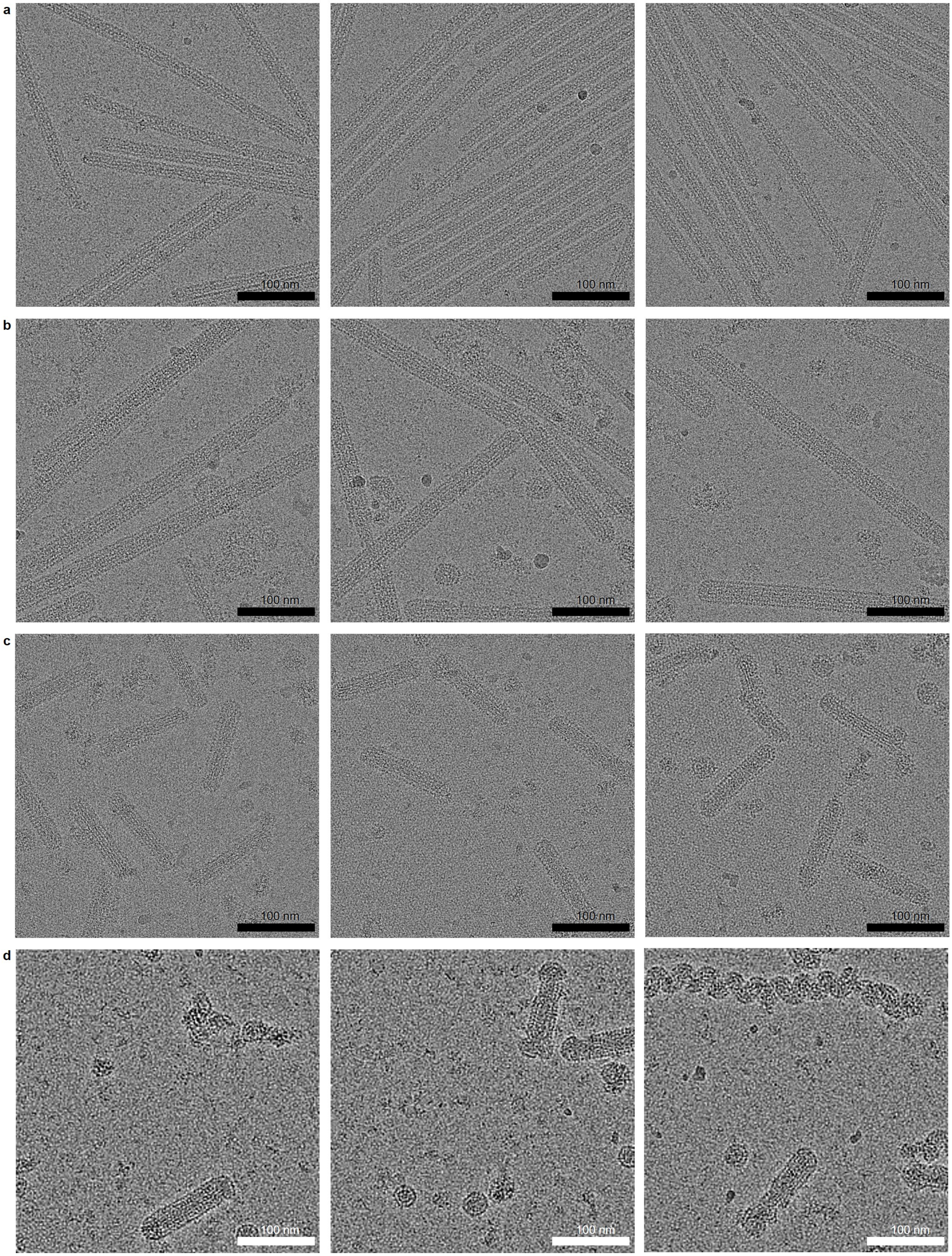


**Fig. S2** Supplementary cryo-EM micrographs for **a**, 6HB-2k, **b**, 6HB-10k, **c**, 24HB-2.5k and **d**, 24HB-10k samples.

#### Note S3: Single-particle reconstruction

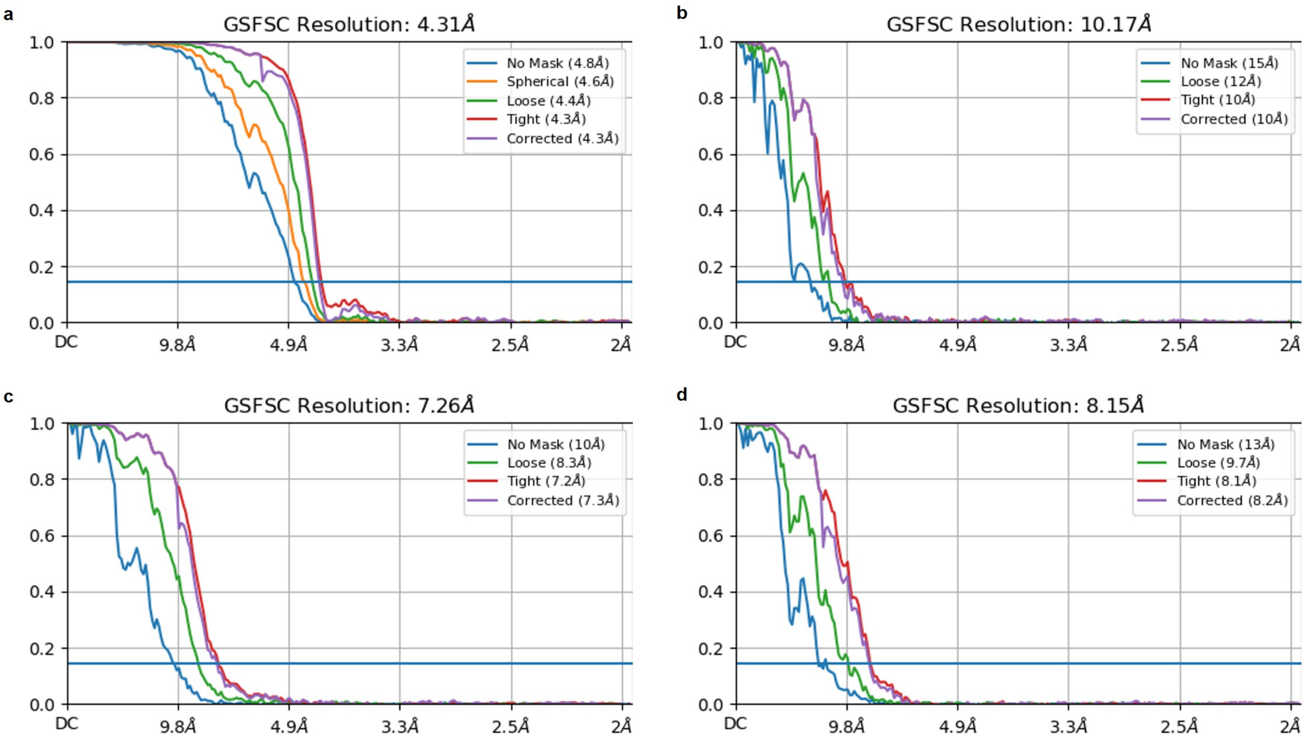


**Fig. S3** Resolution estimates by "gold standard Fourier shell correlation" (GSFSC) for **a**, 6HB-2k, **b**, 24HB-2.5k, **c**, 6HB-10k, first capsid protein (CP) layer and **d**, 6HB-10k, second CP layer.

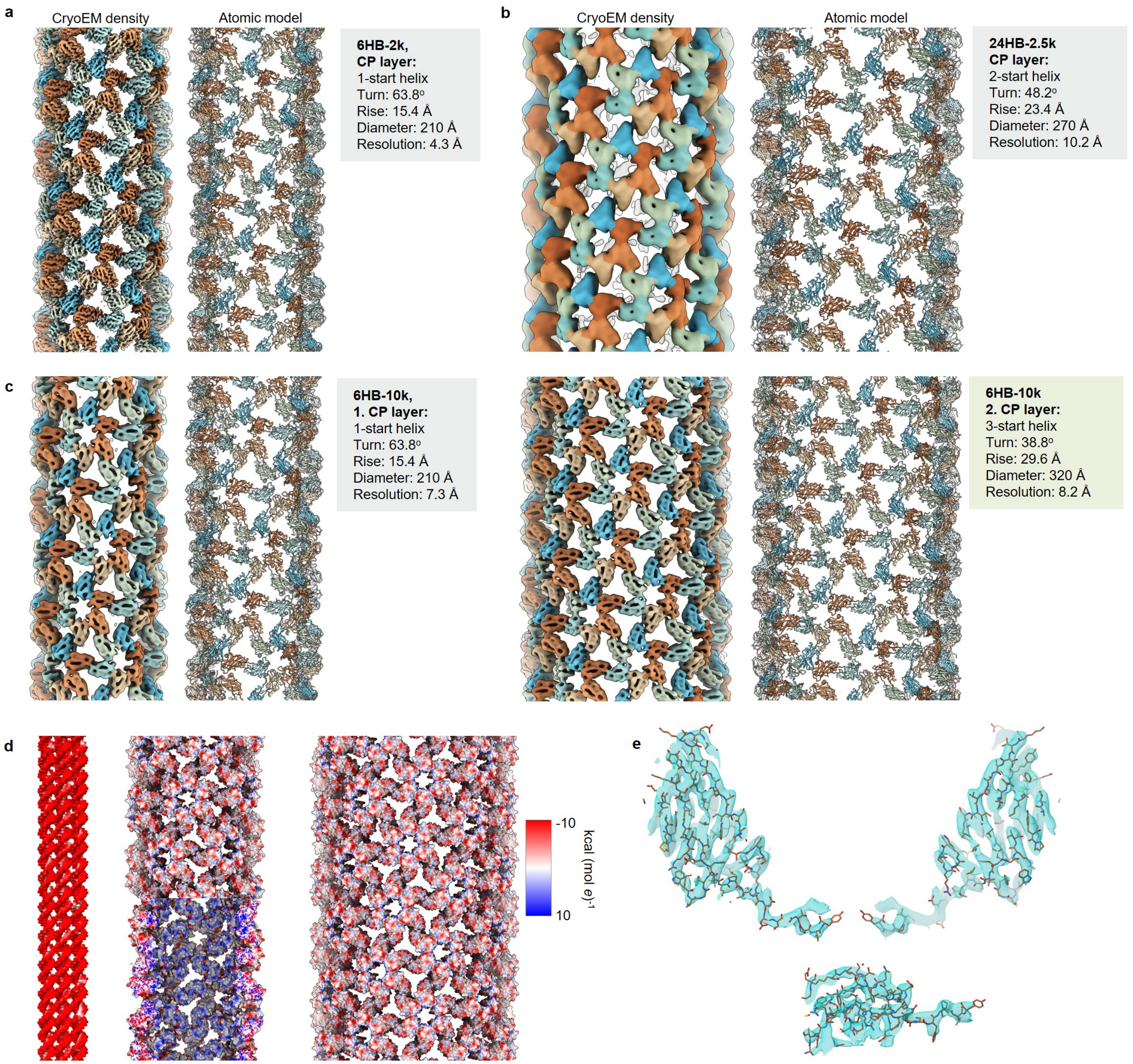


**Fig. S4** Cryo-EM densities (left) and atomic models (right) and characteristics for the CP layers of **a**, 6HB-2k, **b**, 24HB-2.5k and **c**, 6HB-10k. **d**, Electrostatic potential surfaces for 6HB (left), the first CP layer (middle) and the second CP layer (right). **e**, Map to model comparison for the protein shell of 6HB-2k.

#### Note S4: Reconstruction of the cap structure

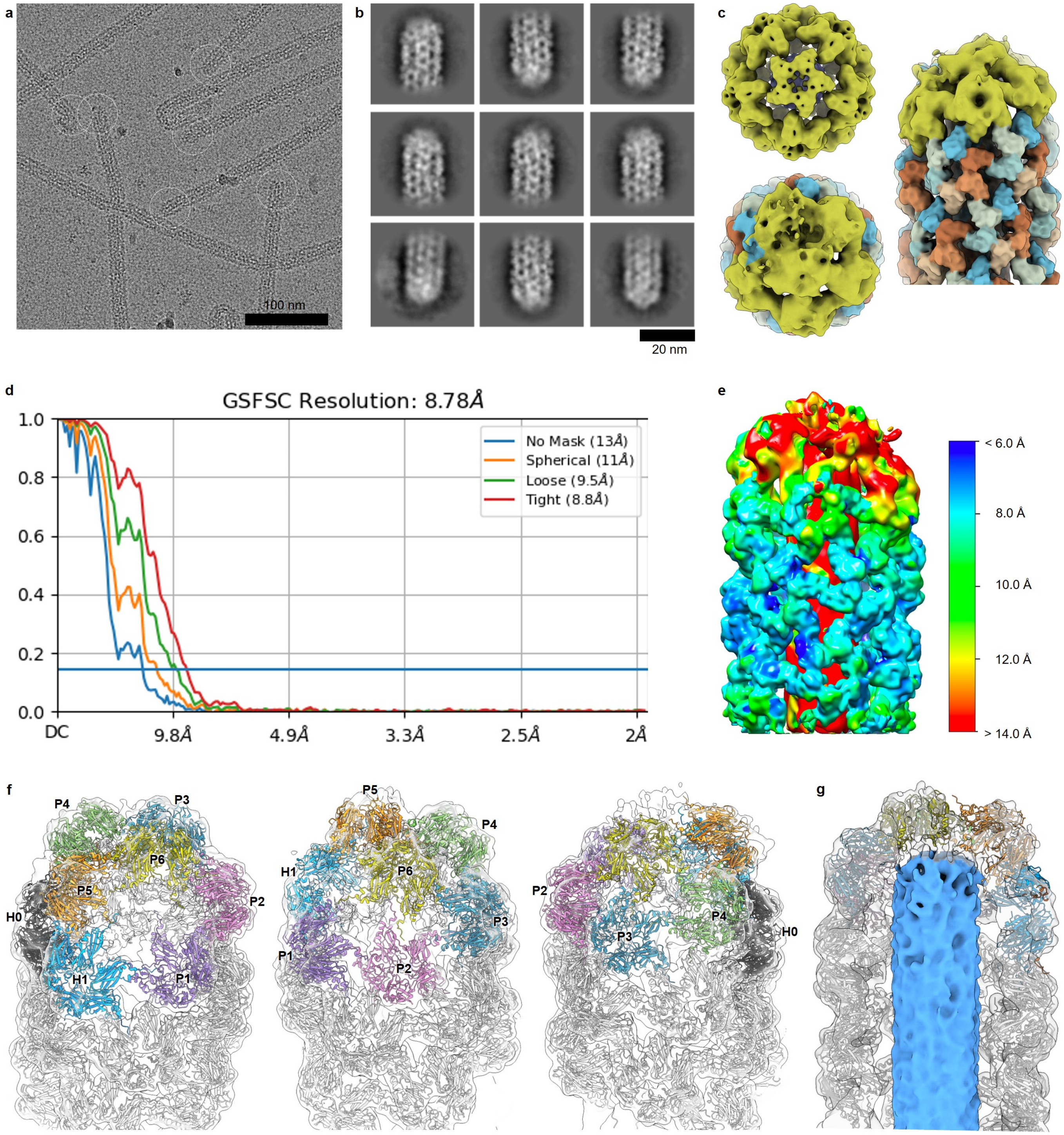


**Fig. S5 a**, Representative cryo-EM micrograph showing the selection the filaments' ends. **b**, Selected 2D class averages. **c**, Comparison between an "empty" spherical particle with $T$ = 1 symmetry (top left; calculated from the 6HB-2k data by icosahedral single particle reconstruction) and the cap structure of 6HB-2k top (bottom left) and side view (right). **d**, Resolution estimates by GSFSC. **e**, Local resolution of the cap structure. **f**, Supplementary views of the cap structure showing the positioning of the hexamers H0-1 (black, blue) and the pentamers P1-6. **g**, Cross-section showing the position of 6HB (blue) in the capped structure.

#### Note S5: Coating of 13HR structure

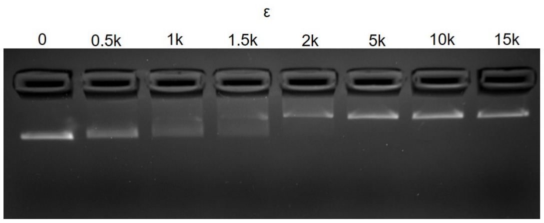


**Fig. S6** The complexation between 13HR and CCMV CPs is monitored using AGE.

#### Note S6: SAXS analysis

For the model, the buffer (see Figure S7a) was subtracted, while a Debye background was added. Additionally, CPs assemble spontaneously into icosahedral structures (1). They can be modelled as solid spheres, resulting in particles with a diameter of ca. 28 nm, suggesting $T = 3$ symmetry (Figure S7b). For 24HB, a cylinder is chosen as its geometrical representative, resulting in a radius of 7.7 nm (Figure S7c). It is notable, that the (following) cylinder models do not include fitting to a Lorentzian peak, which can be used to model the semi-solid intra-bundle distance corresponding to the peak at ca. 0.2 $\text{Å}^{-1}$ (2). As observed in TEM, two different populations, namely the complexed 24HB structures and sphere-like assemblies, are assumed to be present in the complexed sample. Subsequently, the intensity distribution of the capsids is reduced to a form factor by subtraction from the 24HB-2.5k, thus features of the complexed 24HB become visible, and the complexed structures can be modeled as core-shell cylinders (Figure 4d, Figure S7d).

In order to avoid the subtraction factor, the system was furthermore modelled as a three-function model consisting of a core-shell resembling the single protein layer, a cylinder resembling 24HB and a sphere resembling icosahedral CP assemblies (Figure S7e). The model agrees well with the TEM measurements, suggesting diameters of 23.0 nm for 24HB-2.5k and 27.6 nm for spherical assemblies with $T = 3$ symmetry. The thickness of the CP shell on 24HB-2.5k is 4.5 nm. Furthermore, the impact of the single components on the final intensity distribution can be shown with this approach (Figure S7f). The cylinder length, which is expected to be ca. 110 nm, has no impact on the model.

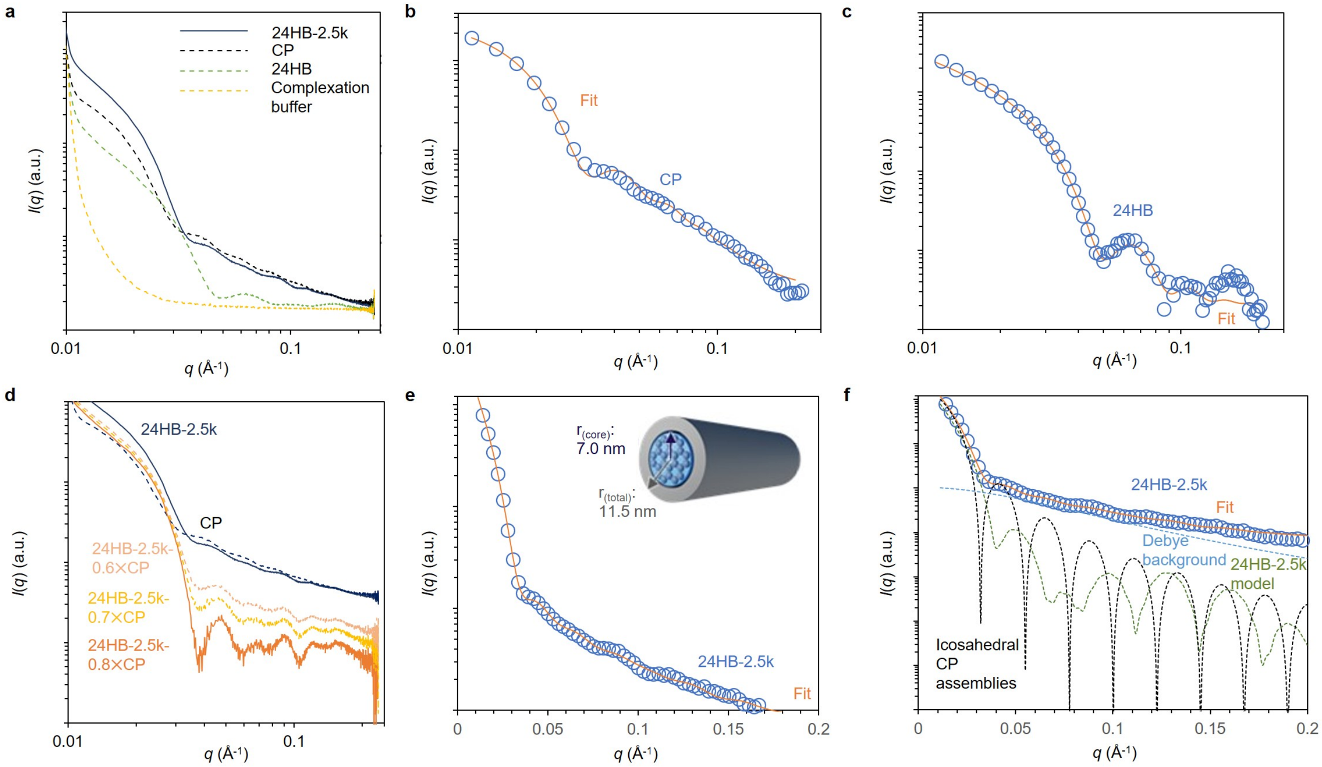


**Fig. S7 a**, Intensity distributions before buffer substraction for 24HB-2.5k (single protein layer, black), CPs only (dark blue dashed), 24HB (green dashed) and the complexation buffer (yellow dashed). The intensity distributions **b**, for sphere-like CP assemblies which are modelled as spheres and **c**, for 24HB a cylinder is used as geometric model. **d**, Influence of the factor used for subtraction of the capsid spectrum from the spectrum of the complexed structure (dark blue) through which features of 24HB become visible. **e**, 24HB-2.5k (blue circles, after background substraction) is modelled (orange) using a three component system consisting of core-shell, cylinder and spheres. **f**, Breakdown of **e** showing the impact of the single components on the final model (orange). Icosahedral CP assemblies (black) are represented by spheres and 24HB-2.5k by a core-shell cylinder (green).

#### Note S7: Negative-stain TEM images

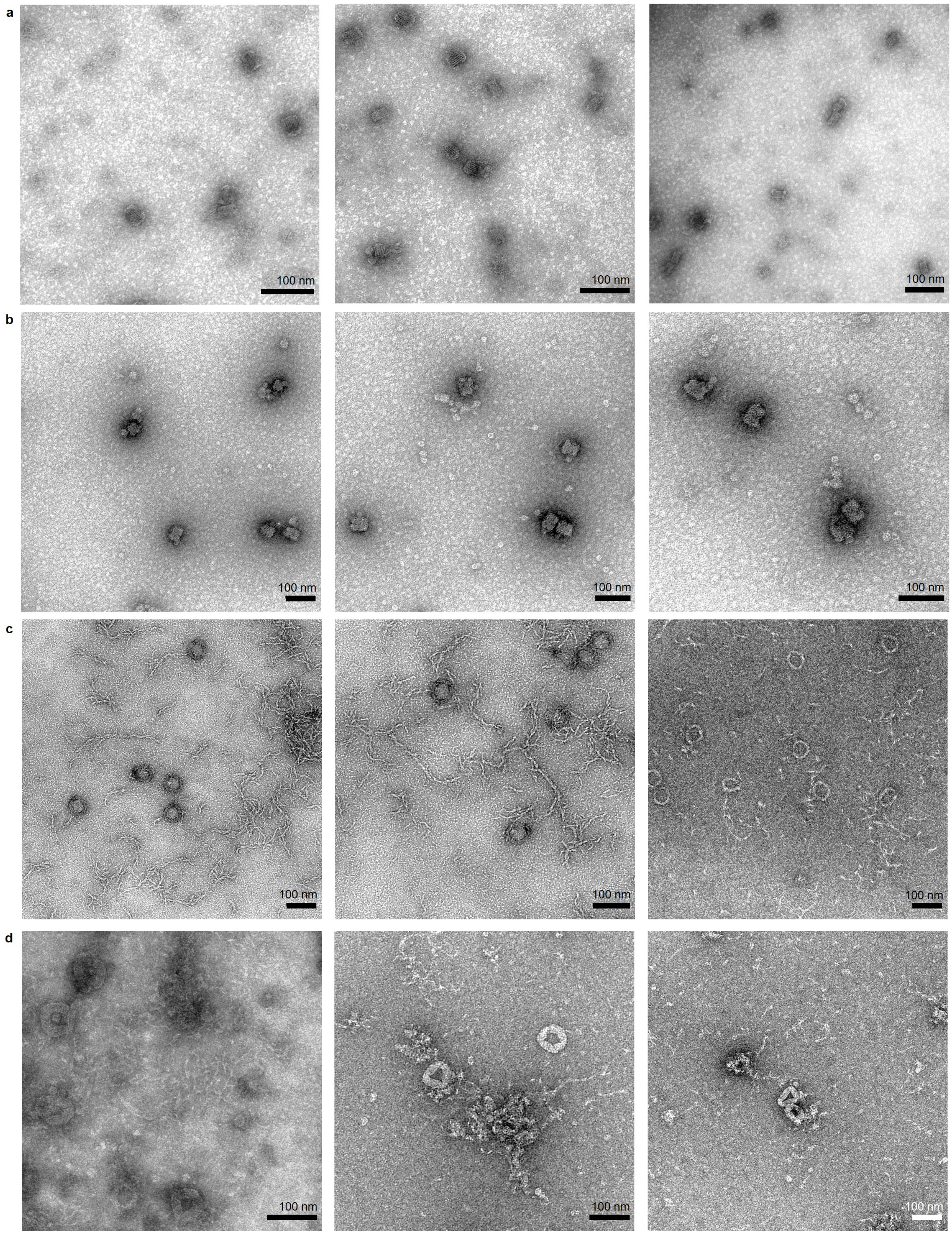


**Fig. S8 TEM images showing origami structures complexed with CPs for full sample representation.** 60HB with **a**, $\varepsilon$ = 2k and **b**, $\varepsilon$ = 10k. 13HR with **c**, $\varepsilon$ = 2k and **d**, $\varepsilon$ = 10k.

#### Note S8: Size distribution of coated structures

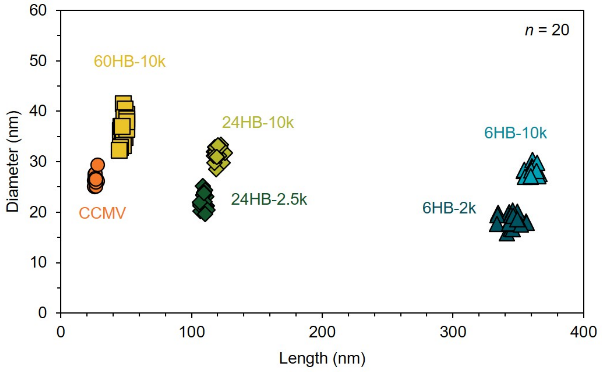


**Fig. S9** Size (length and diameter) of correctly assembled structures showing the uniformity of the dimensions (n = 20). Native CCMV is shown in orange, 6HB samples in shades of blue, 24HB in shades of green and 60HB-10k in yellow.

#### Note S9: Functionalization of 6HB

6HB was functionalized with AuNPs in a two-pot reaction, similar to previously described procedures (3, 4). First, the 6HB structure was prepared as described in the Method section, however, three staple strands were exchanged to contain an overhang which can later hybridize with oligonucleotide-functionalized AuNPs. After purification using poly(ethylene glycol) (PEG) precipitation (as described in Methods, similar to (5)), the folded structures were mixed with oligonuceotide-functionalized AuNPs which were added in 30× excess (10× excess per annealing site), heated to 40 °C and subsequently the temperature was decreased to 20 °C (-0.1 °C $min^{-1}$ ramp).

Briefly, oligonucleotide-functionalized AuNPs were obtained by mixing 40 $\mu$L of AuNPs (5 nm diameter, citrate stabilized, 100 nM, Sigma Aldrich) with 0.8 $\mu$L of sodium dodecyl sulfate (SDS) for 20 min, before incubation with 4 $\mu$L thiol-modified oligonucleotides (for hybridizing with staple overhangs) for 30 min. The AuNPs were salt-aged using 2.5 M NaCl by 6× addition of 0.4 $\mu$L, 6× addition of 0.8 $\mu$L, 5× addition of 1.6 $\mu$L and a final addition of 2 $\mu$L. The interval between the additions is 5 min and all steps are performed at 40 °C and 600 rpm (Eppendorf ThermoMixer C). Subsequently, 60 $\mu$L of 1× folding buffer (FOB, 1× Tris-acetate-EDTA (TAE), 12.5 mM $MgCl_2$) supplemented with 0.02 % SDS are added and the incubation was continued for 1 h before the temperature was decrease to 20 °C for an overnight incubation.

Before usage, the oligonucleotide-functionalized AuNPs were purified from excess oligonucleotides using spin-filtration. After an initial washing step with 200 $\mu$L of the desired buffer (14,000 *g*, 5 min), 360 $\mu$L of AuNPs were added to the filter together with 120 $\mu$L 1× FOB with 0.02 % SDS and centrifuged for 10 min at 14,000 *g*, followed by a 3× addition of 200 $\mu$L of 1× FOB with 0.02 % SDS (10 min, 14,000 *g*). The purified DNA-functionalized AuNPs were recovered by inverting the filter and centrifugation for 2.5 min at 1,000 *g*.

The folded structures were purified from excess staple strands and AuNPs by PEG precipitation (6). Per 50 $\mu$L of folding reaction, 12.5 $\mu$L of PEG buffer containing 17.5 % (w/v) PEG8000, 1× TAE buffer, 500 mM NaCl and 10 mM $MgCl_2$ are used. Before centrifugation at 4 °C and 12,600 *g* for 30 min, the mixture is incubated at 4 °C for 10 min. The supernatant is discarded and the pellet resuspended in 1× FOB with 0.02 % SDS and incubated overnight at 30 °C, 600 rpm on an Eppendorf ThermoMixer C before the procedure is repeated to ensure full removal of excess AuNPs.

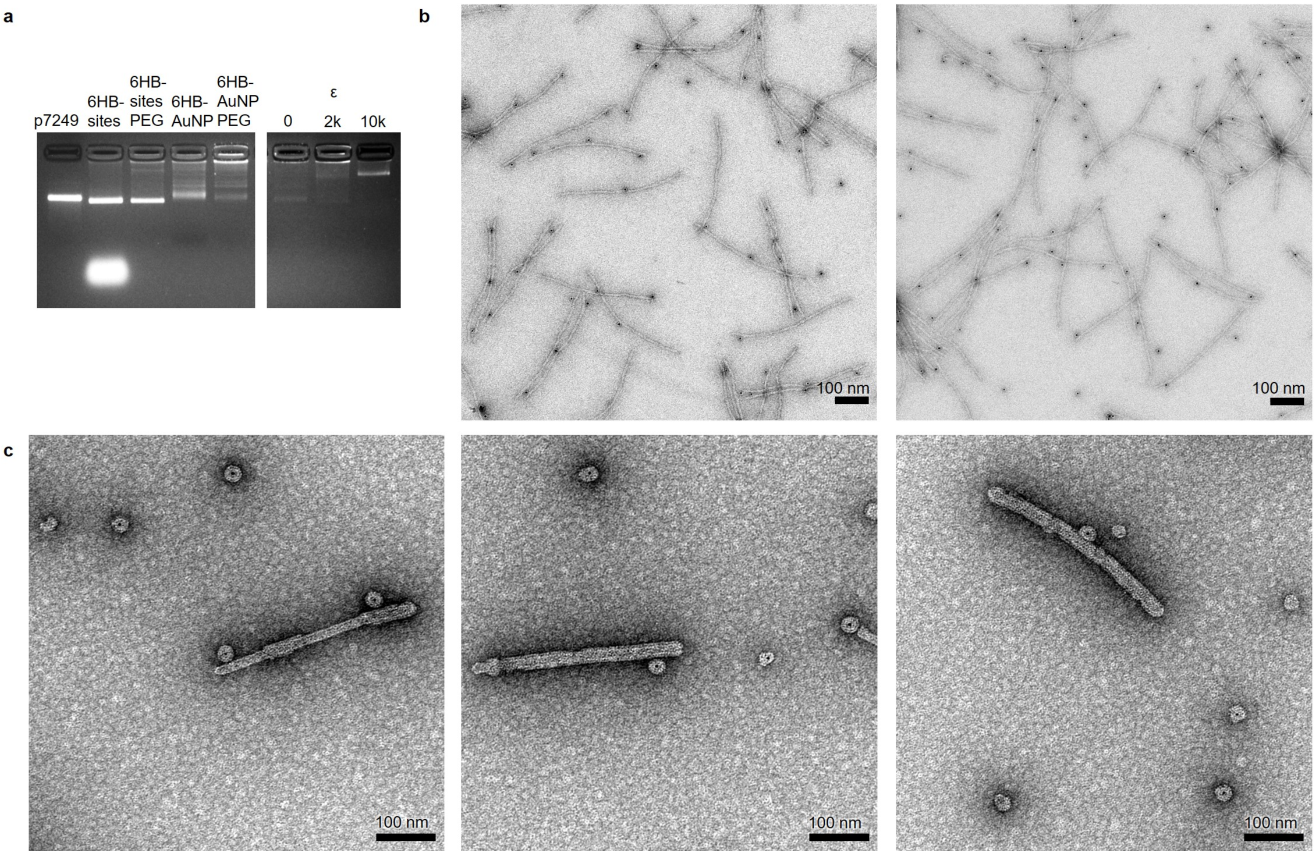


**Fig. S10 a**, Folding and purification of AuNP-functionalized 6HB (left). After purification (lane 3), the staples after the first folding step of 6HB containing AuNP annealing sites (lane 2) have been removed. Subsequently, AuNPs are annealed onto the structure (lane 4, excess gold is represented by the faster migrating, darker band) and in a two-step procedure purified (lane 5). As a reference, the scaffold is shown in lane 1. Complexation of functionalized 6HB with CPs (right). TEM images of **b**, the plain structures in 1× FOB which are buffer-exchanged and coated with CP at **c**, $\varepsilon$ = 10k.

#### Note S10: DNase I digestion studies

Heparin was used as a competitive agent resulting in the disintegration of CPs from the complexed structures resulting in plain origami structures to show the structural intactness after incubation with DNase I. The amount of heparin used is expressed as the ratio between $n_{\text{sulfates}}$ originating from heparin and $n_{\text{phosphates}}$ originating from the DNA backbone. A heparin molecule was estimated to contain on average 71 negatively charged sulfate groups, assuming an average molecular weight of 17–19 kDa and an average of 2.33 sulfate groups per repeating IdoA(2S)-GlcNS(6S) disaccharide unit (7). 6HB origami was estimated to have in total 14569 phosphate groups (3) while 24HB has 15504. For the final digestion experiments, 3.8× excess and 200× excess were used for structures coated with one or two CP layers, respectively.

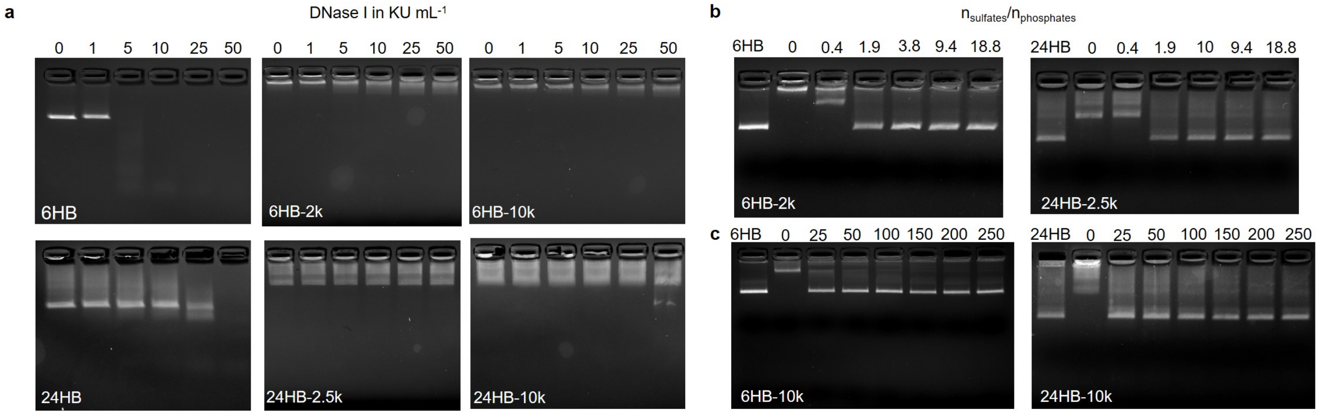


**Fig. S11 a**, Stability against DNase I of 6HB (top) and 24HB (bottom): Plain structures (left) are compared with structures coated with a single (middle) or two protein layers (right) before disassembly of the coating by the addition of heparin. **b**, Excess of heparin required for the release of the 6HB (left) and 24HB (right) when originally complexed with either $\varepsilon$ = 2k or $\varepsilon$ = 2.5k. **c**, Excess of heparin required for the release of the 6HB (left) and 24HB (right) when originally complexed with $\varepsilon$ = 10k.

#### Note S11: Collection of parameters used in cryo-EM and single-particle reconstruction

**Table S1** Cryo-EM structure determination parameters

| | **6HB-2k** (EMD-16076) | **6HB-10k inner** (EMD-16077) | **6HB-10k outer** (EMD-16078) | **6HB-2k cap** (EMD-16079) | **24HB-2.5k** (EMD-16080) |
|---|---|---|---|---|---|
| **Data collection and processing** | | | | | |
| Magnification | 150,000× | 150,000× | 150,000× | 150,000× | 150,000× |
| Voltage (kV) | 200 | 200 | 200 | 200 | 200 |
| Electron exposure ($e^-$/Å$^2$) | 40 | 40 | 40 | 40 | 40 |
| Defocus range ($\mu$m) | 0.7–2.1 | 0.7–2.1 | 0.7–2.1 | 0.7–2.1 | 0.8–2.3 |
| Pixel size (Å) | 0.96 | 0.96 | 0.96 | 0.96 | 0.96 |
| Symmetry imposed | | | | | |
| Point group | C1 | C1 | C1 | C1 | C2 |
| Helical turn (°); rise (Å) | 63.8; 15.4 | 63.8; 15.4 | -107.1; 9.9 | N/A; N/A | 48.2; 23.4 |
| Initial helical segments (no.) | 837,120 | 384,254 | 32,473 | 3904 | 37,075 |
| Final helical segments (no.) | 695,465 | 32,473 | 6141 | 3740 | 4197 |
| Map resolution (Å) | 4.3 | 7.3 | 7.0 | 8.8 | 10.2 |
| FSC threshold | 0.143 | 0.143 | 0.143 | 0.143 | 0.143 |
| Map sharpening *B* factor (Å$^2$) | -300 | -300 | -300 | -300 | -300 |

**Table S2** Model refinement and validation

| | **6HB-2k** (PDB:8BI4) |
|---|---|
| **Refinement & validation** | |
| Model-to-map resolution (Å) | 4.4 |
| FSC threshold | 0.5 |
| Model-to-map CC | |
| Main chain | 0.79 |
| Side chain | 0.78 |
| Model composition | |
| Chains | 6 |
| Non-hydrogen atoms | 6600 |
| Model resolution range (Å) | |
| *B* factors (Å$^2$) | |
| Protein | 34.6 |
| R.m.s deviations | |
| Bond lengths (Å) | 0.004 |
| Bond angles (°) | 1.003 |
| Validation | |
| MolProbity score | 1.29 |
| Clash score | 2.11 |
| Rotamer outliers (%) | 0.14 |
| Ramachandran plot | |
| Favored (%) | 95.63 |
| Allowed (%) | 4.02 |
| Outliers (%) | 0.34 |

#### Supplementary Information references


1. Bancroft, J. B.; Hills, G. J.; Markham, R. A Study of the Self-Assembly Process in a Small Spherical Virus. Formation of Organized Structures from Protein Subunits *in Vitro*. *Virology* **1967**, *31*, 354–379.
2. Fischer, S.; Hartl, C.; Frank, K.; Rädler, J. O.; Liedl, T.; Nickel, B. Shape and Interhelical Spacing of DNA Origami Nanostructures Studied by Small-Angle X-ray Scattering. *Nano Lett.* **2016**, *16*, 4282–4287.
3. Julin, S.; Nonappa,; Shen, B.; Linko, V.; Kostiainen, M. A. DNA-Origami-Templated Growth of Multilamellar Lipid Assemblies. *Angew. Chem. Int. Ed.* **2021**, *60*, 827–833.
4. Ijäs, H.; Hakaste, I.; Shen, B.; Kostiainen, M. A.; Linko, V. Reconfigurable DNA Origami Nanocapsule for pH-Controlled Encapsulation and Display of Cargo. *ACS Nano* **2019**, *5*, 5959–5967.
5. Stahl, E.; Martin, T. G.; Praetorius, F.; Dietz, H. Facile and scalable preparation of pure and dense DNA origami solutions. *Angew. Chem. Int. Ed.* **2014**, *53*, 12735–12740.
6. Shaw, A.; Benson, E.; Högberg, B. Purification of Functionalized DNA Origami Nanostructures. *ACS Nano* **2015**, *9*, 4968–4975.
7. Välimäki, S.; Khakalo, A.; Ora, A.; Johansson, L.-S.; Rojas, O. J.; Kostiainen, M. A. Effect of PEG-PDMAEMA Block Copolymer Architecture on Polyelectrolyte Complex Formation with Heparin. *Biomacromolecules* **2016**, *17*, 2891–2900.
